# The common neural representation in the primary motor area between motor execution and kinesthetic motor imagery

**DOI:** 10.1101/2025.07.11.664262

**Authors:** Fumihito Imai, Jun Shinozaki, Hidekazu Saito, Hiroshi Nagahama, Yuuki Sakurai, Kenji Ogawa

**Author notes:** Correspondence should be addressed to: Dr. Fumihito Imai, Department of Psychology, Hokkaido University, Kita 10, Nishi 7, Kita-ku, Sapporo, 060-0810 Japan, Dr. Kenji Ogawa, Department of Psychology, Hokkaido University, Kita 10, Nishi 7, Kita-ku, Sapporo, 060-0810 Japan.

## Abstract

Although motor imagery activates higher-order motor-related areas, the role of the primary motor area (M1) in motor imagery remains unclear. This study aimed to investigate whether motor imagery recruits a neural representation of fingers similar to that of motor execution in the hand M1. Ten healthy right-handed adults executed and kinesthetically imagined tapping using one of four fingers. Using functional magnetic resonance imaging with multi-voxel pattern analysis, we trained the decoder to classify which finger the participants were moving using brain activation during motor execution and tested whether it could predict which finger the participants were imaging to move during motor imagery (cross-classification). We also performed the classification in the reverse direction. The average accuracy of these cross-classifications was significantly higher than chance in the left hemisphere hand M1 (hand-M1). Analysis of the representational geometry showed that the distance of neural representations for the same fingers was statistically shorter than that for different fingers between motor execution and imagery. Furthermore, we conducted a replication study with 14 participants and found results similar to those of the original study. Our results suggest that the neural representation of kinesthetic motor imagery is partially similar to that of motor execution in the contralateral hand M1.

## Introduction

Motor imagery is defined as the “mental simulation” or “mental rehearsal” of motor activity without muscle activity (Decety, 1996; Grush, 2004; see also Jeannerod, 1994). Although motor imagery activates higher-order motor-related areas, the role of the primary motor cortex (M1) in motor imagery is unclear (Hanakawa, 2016; Sharma and Baron, 2014). Hétu et al. (2013) reported that only 18% of 122 studies detected M1 activation during motor imagery (see also Lebon et al. 2018; Naito et al., 2002; Porro et al., 1996; Roland et al., 1980; Shibasaki et al., 1993). A review by Hardwick et al. (2018) concluded that the relationship between motor imagery and M1 was not supported by meta-analysis. Although some researchers have discussed the possibility that the M1 encodes motor direction during motor imagery (Lotze and Zentgraf, 2010) or that the M1 is related to motor imagery at the motor preparation stage rather than at the planning stage (Hanakawa, 2016), there is little evidence that the M1 is related to motor imagery. Using multi-voxel pattern analysis (MVPA), Zabicki et al. (2017) and Monaco et al. (2020) found no common neural representation between motor execution and imagery in the M1.

Why is M1 activation unstable during motor imagery? One possible cause for this discrepancy is that imagery contains various aspects (Hanakawa, 2016; Lotze and Zentgraf, 2010). According to Hanakawa (2016), motor imagery typically involves two aspects: visual and kinesthetic. The visual aspect (i.e., visual motor imagery) is the visualization of motor action, while the kinesthetic aspect (i.e., kinesthetic motor imagery) is the kinesthetic feeling of body movements without executing movements (e.g., Guillot et al., 2009). Previous studies have suggested that visual motor imagery mainly depends on visual-related areas and the superior parietal lobule, whereas kinesthetic motor imagery evokes the activation of motor-associated structures, including the M1 and the inferior parietal lobule (Guillot et al., 2009; Schnitzler et al., 1997; see also Stinear et al., 2006). Another factor is the complexity of imagined movements. Previous research (Zabicki et al., 2017) investigated the presence of shared neural representations in motor execution and imagery for right-hand aiming, extension-flexion, and squeezing and found no such representations in M1 (see also Monaco et al., 2020). This may be due to a combination of factors related to movement and effectors. Therefore, the present study aimed to eliminate these confounding factors by using simple repetitive finger tapping, focusing solely on the difference in effectors.

The present study aimed to test whether motor imagery recruits similar neural representations of the fingers to those of motor execution in the M1. The participants executed or kinesthetically imagined tapping with one of their right fingers. Using functional magnetic resonance imaging (fMRI) with MVPA, we attempted to classify the content of kinesthetic motor imagery with brain activation based on those of execution, and vice versa (cross-classification: Kaplan et al., 2015). If motor execution and kinesthetic motor imagery share a common neural representation, cross-classification between motor execution and kinesthetic motor imagery would be successful. To determine the extent to which neural representations were similar between motor execution and imagery, we conducted a representational similarity analysis (RSA; Kriegeskorte et al., 2008). Using RSA for simple finger movement execution, Ogawa et al. (2019) found that two adjacent fingers had more similar neural representations than more distant fingers. Ejaz et al. (2015) reported that the neural representation of the middle and ring finger pair or the ring and little finger pair was highly similar to that of the other finger pairs. If kinesthetic motor imagery has a representational geometry similar to that of motor execution, we might find the same effects reported by Ejaz et al. (2015) and Ogawa et al. (2019) for not only executed but also imagined finger movements. Another prediction was that if the representational geometry of fingers is similar between motor execution and imagery, the distance of neural representational similarity between motor execution and imagery should be shorter for the same finger pairs than for different finger pairs. To verify these predictions, we implemented RSA, including the representational dissimilarity matrix (RDM) and a classical multi-scaling method.

## Materials and methods

### Participants

Ten males with normal or corrected-to-normal vision and no neural disease participated in the experiment (mean age, 27.2 years; range, 23–38 years). All participants were right-handed as defined by a modified version of the Edinburgh Handedness Inventory (Oldfield, 1971) for Japanese individuals (Hatta and Nakatsuka, 1975). They had normal or corrected-to-normal vision and no neural disease. All participants provided written informed consent before the experiment in accordance with the Declaration of Helsinki. The local ethics committee approved the experimental protocol.

### Task procedures

First, the participants completed the Movement Imagery Questionnaire-Revised (MIQ-R; Hall and Martin, 1997) for Japanese (JMIQ-R; Hasegawa, 2004). They executed and then imagined the instructed action, either visualizing the action from a third-person perspective (third-person perspective visual motor imagery) or kinesthetically feeling the body movement while visualizing the action from a first-person perspective (performer’s own perspective: kinesthetic motor imagery + first-person perspective visual motor imagery). There was a total of eight trials as two imagery modalities by four actions: leg movement, jumping, arm movement, and trunk flexion. After each instance of imagery, the participants rated how easy it was to visualize (third-person perspective visual motor imagery) or feel (kinesthetic motor imagery + first-person perspective visual motor imagery) the action on a 7-point Likert scale: 1 was very hard and 7 was very easy. This questionnaire was administered prior to the MRI sessions on the same day or within 1 week prior to MRI.

The participants then underwent three types of MRI scans: a fixation session, a functional localizer scan, and subsequent execution and imagery task sessions, four each for execution and imagery in that order. The stimuli were presented using PsychoPy (version 1.83.04; Peirce, 2009, 2007).

To rule out the possibility of participants moving their fingers while imagining finger movements (Hanakawa, 2016), we simultaneously performed MRI and electromyography (EMG). Participants underwent a fixation session by lying in the MRI scanner with an EMG apparatus placed on the hypothenar, extensor digitorum communis, extensor indicis proprius, and flexor digitorum superficialis muscles of both hands and forearms. They saw the fixation cross in white on a gray background displayed at the center of a nonmagnetic screen through head-mounted mirrored goggles. They rested while looking at the central fixation cross for five minutes, which was the same length as the session for the following execution/imagery task sessions. The EMG data extracted from this session were used for the rest period in the EMG analysis (see details in Supplementary EMG recording and processing). We used the MRI scanner to record the EMG of a rest period in the same situation as the later execution/imagery task session, i.e., scanner sound, potential MRI scan noise in the EMG data. Therefore, the MRI data from this session were not used in the analysis.

The next functional localizer scan was conducted to identify each participant’s activated voxels related to finger movements. First, only the center fixation cross was displayed for six seconds, and then the word “start” appeared above the center fixation cross for three seconds. Subsequently, task and rest blocks were initiated. The task and rest blocks alternated 12 times. Each task block began when the word “MOVE” was displayed above the central fixation cross. Simultaneously, a countdown number from 12 to 1, representing seconds, was displayed below the center fixation cross. The participants executed 12 repetitive movements of opening and clenching their right hand, synchronized with the displayed number. Then, the word “MOVE” disappeared and a new countdown from 12 to 1 appeared below the fixation cross, indicating the start of a rest block. The participants did not move any parts of their bodies during this block. All stimuli were shown in yellow during the task block and white during the other periods. The functional localizer session required five minutes.

Four execution task sessions and four subsequent imagery task sessions were conducted after the functional localizer session. The participants held a four-button response pad (Current Designs, USA.) in each hand; the right one was for performing the execution task, and the left one was for self-evaluation of the execution and imagery tasks. Each button on the right response pad corresponded to the right index to the little finger. In the first four sessions, the participants executed the 1-Hz tapping of one of the four right fingers. In the last four sessions, participants imagined the same one-finger tapping. This order allowed participants to become familiar with the kinesthetic sensation of finger tapping that they imagined during the imagery task sessions. Each session consisted of 12 task blocks, each followed by a 12-second non-task period. Each period was divided into a five-second evaluation, a five-second rest period, and a two-second instruction for the next task. Each instance of finger tapping was executed and imagined in three blocks in each session. Thus, there were three blocks × four sessions for a total of twelve task blocks for each task condition of the four fingers by two modalities. The order of the required finger tapping was randomized for each session. Details of the imagery instruction were as follows: “Imagine kinesthetically the tapping of your fingers in sync with a countdown that is the same as the execution task. You can feel your movement internally, even if your eyes are closed. Kinesthetic imagery is similar to this sensation”. The stimulus presentation was identical to that described by Ogawa et al. (2019). First, a white Japanese character appearing above the central fixation cross indicated the finger to be moved or imagined to be moved. Two seconds later, a countdown number from 12 to 1 for each second was displayed below the center fixation cross. In the respective task session, the participants executed or imagined 12 repetitive tapping movements with their required finger, synchronized with the displayed number. Participants had to look at the display and keep their eyes open to synchronize with the countdown, regardless of the execution or imagery task. For the subsequent five seconds, a Japanese word meaning “evaluation” and four options listed in Japanese, meaning “A. Very easy,” “B. Easy,” “C. Difficult,” and “D. Very Difficult,” appeared in tandem. Participants evaluated the quality of their last action in the execution task block and the ease of their last imagined action in the imagery task block. They then pressed the corresponding button using their left index finger. For the next five seconds, a fixation cross appeared at the center of the screen, and a countdown number from five to one appeared below it. The participants remained still for those five seconds (Figure 1 top). Any kind of button pressing was recorded. In the execution task, the error block was defined as the block in which a participant pressed the button differently than instructed for more than half of the total taps in that block; for example, 6 out of 12 taps in a block.

**Figure 1:**
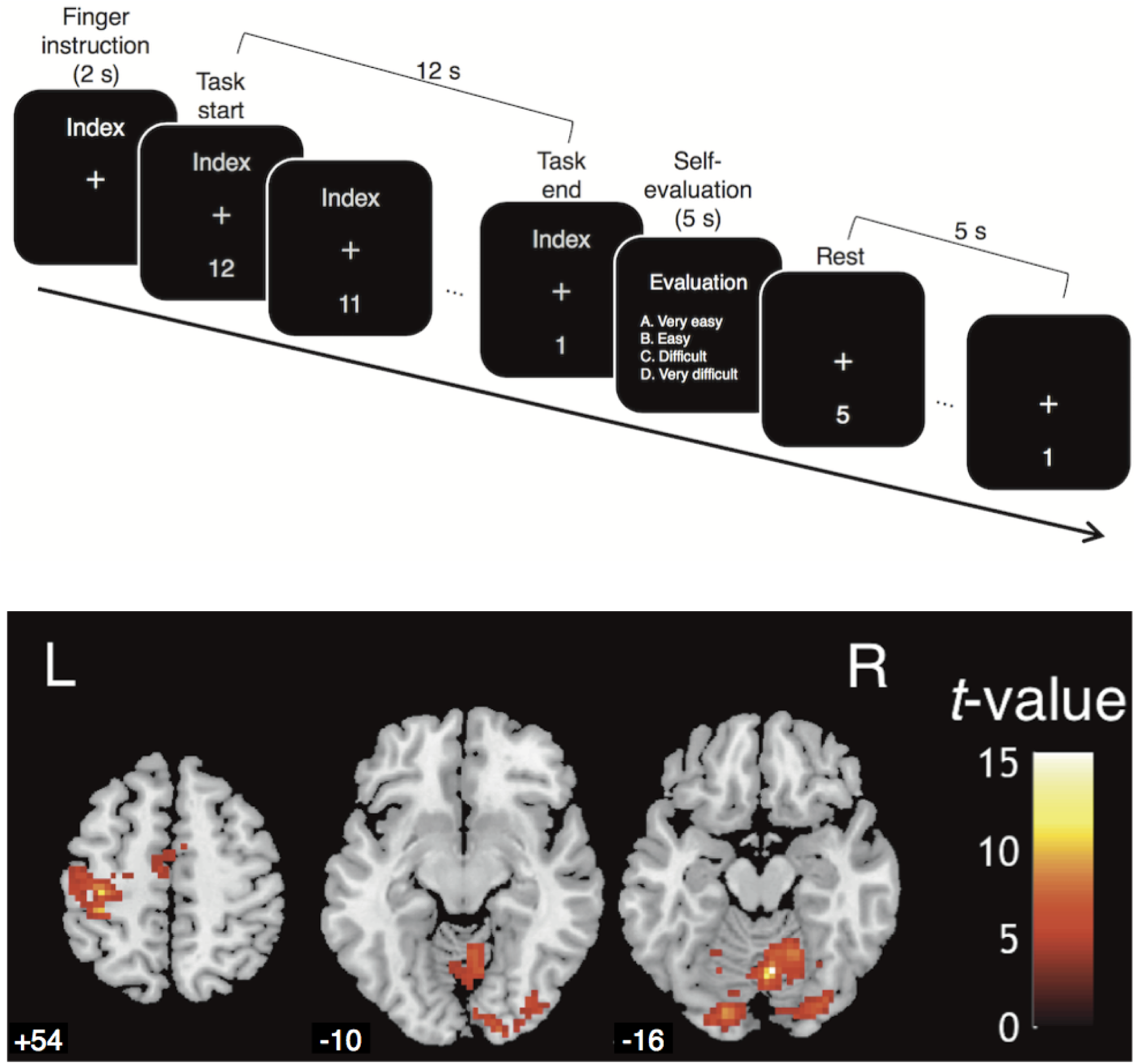
Top) The timeline of a task and rest block in the execution and imagery task sessions. Although English words appear in this Figure, Japanese characters have been used in the experiment. Bottom) Areas activated during the functional localizer scan based on the group-level with a threshold of *p* < 0.001, k ≥ 50, *n* = 10. The three brain images represent slices of the axial view at Z = +54 as sensorimotor areas and Z = −10 and −16 as visual areas, respectively. Note that the actual mask for each participant’s ROIs was created using the individual results with a threshold of *p* < 0.01, k ≥ 0. L indicates the left hemisphere, and R indicates the right hemisphere.

### Experimental design and statistical analyses

The experimental design had a two-factorial within-subject setup: task (motor execution or motor imagery) × finger (right index finger to the little finger). We also used a two-factorial within-subject design for the EMG data: task (motor execution, motor imagery, or rest) × side (left or right forearm). We used a two-way within-subjects analysis of variance (ANOVA) to analyze the EMG data and test for interactions between the side and task. If our task worked correctly, a large EMG activity would only be detected on the right hand while performing the execution task. We also used a two-way within-subjects ANOVA for RSA to test whether there were main effects of task condition (execution vs. imagery) and finger pair (six finger pairs) without interaction. A one-way within-subjects ANOVA was performed on the behavioral data (i.e., time-gap performance in the execution task) to test whether performance differed between fingers. Two-tailed one-sample *t*-tests were adapted for MVPA decoding accuracy to examine whether the decoding accuracy was above chance level. In the RSA, two-tailed paired *t*-tests were used to compare the distance between neural representations: the mean distance between the executed and imagined same finger movement pairs versus the mean distance between the executed and imagined different finger movement pairs. To correct for multiple comparisons, we utilized the Holm–Bonferroni procedure (Holm, 1979) to analyze the MRI and behavioral data and Tukey’s method for the EMG data analysis. The statistical significance level for each analysis is reported in the following section.

### MRI acquisition

All MRI was conducted at Sapporo Medical University Hospital using a Philips Healthcare (Amsterdam, Nederland) 3-Tesla MRI scanner with a 32-channel head coil. T2*-weighted echo-planar imaging (EPI) was used to acquire 84 scans for the localizer and 100 scans for each fixation session and execution/imagery task session using a gradient-echo EPI sequence. The first two scans from each session were discarded to allow the magnetization to reach a stable state. The scanning parameters of fMRI were as follows: repetition time (TR), 3000 ms; echo time (TE), 30 ms; flip angle (FA), 90°; field of view (FOV), 192 × 192 mm; matrix, 96 × 96; 50 axial slices; and slice thickness, 3.0 mm without a gap. T1-weighted anatomical imaging was performed with the following parameters: TR, 5.871 ms; TE, 3.454 ms; FA, 13°; FOV, 256 × 256 mm; matrix, 256 × 256; 1 mm-cubic 3D scan.

### Processing of fMRI data

Image processing was performed using SPM12 (Wellcome Department of Cognitive Neurology, http://www.fil.ion.ucl.ac.uk/spm). First, all functional images were realigned for motion-related artifact cancellation and then spatially normalized with the Montreal Neurological Institute (MNI) template based on the affine and nonlinear registration of co-registered T1-weighted anatomical images (normalization procedure of SPM). Second, we resampled the normalized images into 3 × 3 × 3 mm voxels with sinc interpolation and spatially smoothed the images using a Gaussian kernel with a 6 × 6 × 6 mm full width at half-maximum. We conducted smoothing only to the mass-univariate analysis to avoid blurring fine-grained information of the multi-voxel activity (Kamitani and Sawahata, 2010; Mur et al., 2009). Next, based on the general linear model, we processed 12 blocks of each session by 12 independent boxcar regressors convolved with a canonical hemodynamic response function. Then we applied a high-pass filter whose cutoff period was 128 seconds to remove low-frequency noise. Finally, serial correlations among the scans were estimated using an autoregressive model. As a result, beta values were obtained as 12 independently estimated parameters of each voxel for respective sessions.

### Definition of regions of interest (ROIs)

The region of interest (ROI) was defined using a combination of anatomical and functional ROIs. The anatomical ROIs were defined as follows: The left M1, bilateral supplementary motor area (SMA), and visual area as Brodmann areas (BA) 17 and 18 were defined using the automated anatomical labeling (AAL) toolbox (Tzourio-Mazoyer et al., 2002). The left M1 and bilateral SMA were masked using the Human Motor Area Template (HMAT: Laboratory for Rehabilitation Neuroscience, http://lrnlab.org/).

Functional ROIs were defined at the individual level based on the functional localizer session. We searched for areas activated during right-hand opening and squeezing movements using the contrast image of the task block versus the implicit baseline for each participant, calculated with a fixed-effects model (see also Figure 1 bottom and Supplementary Table 1). We analyzed the contrast images based on a random-effects model using a one-sample *t*-test for each participant. A significance threshold was set at *p* < 0.01 uncorrected, k ≥ 0. For individual participants, the obtained significant voxels were used as a mask for anatomical ROIs.

This masking process resulted in three ROIs for each participant: hand M1 of the left hemisphere (left hand-M1), bilateral SMA, and bilateral visual area. The mean voxel numbers (SDs) of these ROIs were 370.5 (90.3), 127.2 (66.3), and 300.6 (159.4), respectively. Besides the left hand-M1, the bilateral SMA was chosen because previous studies consistently reported associations with motor imagery (e.g., Hanakawa, 2016; Park et al., 2015). The bilateral visual area was used as a control ROI because the total voxel numbers of BA 17 and 18 were comparable with those of the left hand-M1. We also selected the cerebral white matter and cerebrospinal fluid (CSF) as other control ROIs. We created these ROIs by reducing the above 3 ROIs (M1, SMA, and visual area) from the cerebral white matter and CSF of the WFU_pickatlas (Maldjian et al., 2003) for each participant.

### fMRI mass-univariate analysis

Conventional mass-univariate analysis of individual voxels was performed to identify the activated areas for motor execution and motor imagery. We compared activity during execution/imagery task blocks with that of the implicit baseline in each ROI. In addition, we compared the activity between execution/imagery task blocks and rest blocks. A rest block was defined as the five seconds after a task block, during which participants were required not to make any movements (see Task procedure). We created contrast images for each participant using a fixed-effects model and analyzed the beta values of these images using a random-effects model of a one-sample *t*-test.

We calculated the mean signal change rate of individual ROIs based on the beta values of each voxel and estimated the activation changes in the execution/imagery task blocks. For the respective task sessions with the four-finger collapse, a two-tailed *t*-test was used to determine whether the activation change of the task blocks differed from no change, i.e., an implicit baseline. We corrected the significance level for multiple comparisons using the Holm–Bonferroni procedure (Holm, 1979).

### Multi-voxel pattern classification

Multi-variate pattern analysis was conducted to classify the finger that was executed or imagined for tapping. We attempted to classify using a multi-classifier based on a linear support vector machine using LIBSVM (http://www.csie.ntu.edu.tw/~cjlin/libsvm/) with a fixed regularization parameter of C = 1. The inputs to the classifier were parameter estimates, i.e., beta values without normalization. For all types of classifications, we attempted to predict which finger the participants were moving or imagining to move by the “one-against-one.” It made *k* (*k* - 1)/2 binary classifiers (*k* = number of the task conditions: e.g., four fingers in the within-classification). Each binary classifier processed one of all the pairwise comparisons between the fingers and voted on which of the two candidate fingers better matched the input dataset, i.e., the parameter estimates (beta values). Finally, the finger with the highest number of votes was adopted as the label for the given datasets. It was theoretically possible that one label was not accepted even once, whereas the other was accepted more than four times.

The averaged classification accuracy of within classification for each of the execution and imagery tasks was estimated by a fourfold “leave-one-out” cross-validation. The classifier was trained and tested using datasets from either the execution or imagery sessions. The parameter estimates were beta values from three of the four same-task sessions for the training data and from the remaining session for the test data.

In the cross-classification, the classifier was trained to accurately assign one of the four fingers as the condition label to the input dataset obtained from the execution task sessions. It was then tested to determine whether it could correctly classify each block of the imagery task session dataset into one of the four imagined finger movements. The reverse direction classification was also performed. Training was performed using all data from one of the two tasks, and testing was performed using all data from the other task. Finally, the average cross-classification accuracy score was calculated based on the scores obtained by these two cross-classification directions.

Even if cross-classification between executed and imagined movements is successful, there should be some differences between motor execution and imagery. To detect such a difference, we attempted to distinguish between imagined and executed movements using a between-modality classification. This classification was performed by a fourfold “leave-one-out” cross-validation. For each fold, three of the four sessions from both the execution and imagery tasks (e.g., sessions one through three) were used for training, and the remaining session from the execution task (e.g., session four) was used for testing. The reverse direction, in which the classifier was tested with the remaining imagery task session, was also performed. The final classification accuracy between the modalities was calculated by averaging the accuracies between each fold and the classification direction.

For the ROI analysis of MVPA, we attempted three types of classification: two within-classifications and one cross-classification. The resulting three classification accuracies from within-execution, within-imagery, and the average of the cross-classifications were compared to the chance-level (25%) using two-tailed *t*-tests, with inter-subject differences as random factors and degrees of freedom = 9. Permutation tests were also performed using the same computations as in the above classifications, except for one difference: the classifier for the within-execution/imagery classifications trained on the correct datasets was tested on the datasets in which the finger labels and parameter estimates were shuffled. The cross-classification classifier was tested on datasets in which the finger and modality labels and parameter estimates were shuffled. We repeated this process 1000 times separately for each ROI, shuffling the labels each time, and obtained an empirical distribution of the classification accuracy under the null hypothesis of chance. If the classification accuracy with correct labeling exceeded the 95th percentile of this null distribution, it was judged to be significantly above chance accuracy at *p* < 0.05. In all cases, the significance level for multiple comparisons was corrected using the Holm– Bonferroni procedure (Holm, 1979).

The searchlight analysis (Kriegeskorte et al., 2006) was similar to the ROI analysis, except that the multi-voxel activation patterns of the area within the searchlight spot were targeted instead of an ROI. The searchlight spot was a 9 mm-radius sphere consisting of 100 to 123 voxels as the minimum-to-maximum values. The searchlight spot effectively moved the gray matter of the entire brain. For within-execution/imagery classification, the average classification accuracy with leave-one-session-out cross-validation was allocated to the center voxel of the sphere for each searchlight spot. The method of statistical analysis was the same as that for the ROI analysis, except that permutation tests were not performed because of the time required. For the two within-classifications (execution and imagery) and cross-classifications, activation was reported with a voxel-level threshold of *p* < 0.0005 uncorrected for multiple comparisons with an extent threshold of k ≥ 10.

### Representational Similarity Analysis

We performed RSA to examine the representational geometry for each pair in the executed and imagined tasks: (four fingers in the execution task + four fingers in the imagery task) × (four fingers in the execution task + four fingers in the imagery task) = 64 finger pairs. The RDM was computed based on the same parameter estimates, i.e., the beta value, as the classification analysis. The measure of dissimilarity was the cross-validated Mahalanobis distance (Ejaz et al., 2015; Ogawa et al., 2019) calculated at the individual level. As a “leave-one-session-out” method, one session of each task (execution and imagery) was assigned to one dataset. Sessions with the same number were paired; for example, the first sessions of the execution and imagery tasks were paired. The remaining six sessions, consisting of three sessions of execution task + three sessions of imagery task, were assigned to the other dataset. Thus, there were four folds, and the distance values were averaged across all folds. To calculate the cross-validated Mahalanobis distance, we relied on the optimal shrinkage algorithm (Ledoit and Wolf, 2003) to estimate the voxel-by-voxel noise covariance matrix for each dataset, which was also used by Ejaz et al. and Ogawa et al. However, in only a few cases did the estimated covariance matrix become hardly invertible, and the subsequent computation became unstable, i.e., MATLAB warned that the matrix was close to singular or badly scaled. As a solution, we used 0.4 as the shrinkage coefficient applied to the estimation of the voxel-by-voxel noise covariance matrix because this value usually works well for fMRI data (Diedrichsen et al., 2016). The off-diagonal elements of the RDM indicate the difference in activation patterns between the executed/imagined finger movements.

### Replication study

Because the sample size of the first study was relatively small, we conducted a replication study to assess the reproducibility of our original results, particularly cross-classification accuracy. Sixteen males with normal or corrected-to-normal vision and no neural disease participated in this study (mean age, 23.0 years; range, 20–27 years). None of the participants had participated in the first study. Two participants were excluded from analysis. One exhibited a large head motion: more than 2 mm translation or framewise displacement eight times across all task sessions. The other did not respond to the questions in the eight blocks during one imagery task session. According to G*Power 3.1.9.2 (Faul et al., 2009, 2007), based on the effect size of the cross-classification decoding accuracy obtained in the first study, to obtain a statistical power of 0.8 or higher, the required minimum sample size was 11 participants. Thus, data from the 14 participants in this study achieved sufficient statistical power.

The task procedures and analyses used were the same as those in the first study, except that imagery questionnaires and EMG recordings were not performed. Imagery questionnaires were omitted because we were able to compare the imagery abilities of the participants in the original and replication studies based on their responses regarding the ease of kinesthetic motor imagery obtained during MRI sessions of the imagery task.

All MRI was performed using a Siemens (Erlangen, Germany) 3-Tesla Prisma scanner with a 64-channel head coil at the Center for Experimental Research in Social Sciences, Hokkaido University. The number of scans and scanning parameters were the same as those for the first study, except that, to ensure that the spatial resolution was set to the same as that of the first study, the TR was 3050 ms. If we obtained similar results of the cross-classification to those of the first study using a different MRI and the group of participants, this would indicate the robustness of our main findings.

## Results

### Behavioral analysis

The mean JMIQ-R scores (SDs) for all participants were 5.525 (1.341) for kinesthetic, 5.275 (0.961) for visual, and 5.400 (0.620) for both combined. These scores were significantly higher than the median score of 4 on a 7-point Likert scale, as shown by a one-sample two-tailed *t*-test (*t*s (9) > 3.597, *p*s < 0.01) with a threshold *p* < 0.05, corrected using the Holm–Bonferroni method. This indicated that the participants found the imagery of movements relatively easy. Therefore, we concluded that the participants had sufficient imagery abilities to participate in the experiment. There were only three error blocks out of the 480 execution task blocks for all participants, consisting of one error block for one participant and two error blocks for another participant. Therefore, we included data from all 10 participants in the analysis.

We also compared the tapping performances of different fingers during the execution task sessions. The “time gap performance” was calculated as follows: We averaged the time difference between the appearance of countdown number 12 and the first button tap of the task block, and also the time difference between each successive tap. These calculations excluded incorrect button taps, i.e. pressing a button that did not correspond to the instructed finger. The average temporal difference between taps was calculated for the respective fingers using these time differences. The time gap performances of all fingers were marked less than 1 s. This might be because the time difference between the appearance of the countdown number 12 and the first tap, as well as that between the first and second taps, was typically less than 1 s. Finally, we compared the average time difference between fingers using a one-way within-participant ANOVA (four levels: index, middle, ring, and little fingers). The mean (SD) values for the fingers were 0.925 (0.024), 0.929 (0.021), 0.92 (0.016), and 0.926 (0.017), respectively. The main effect of the finger was not significant (*F* (3, 27) = 1.297, *_p_*η*^2^* = 0.126, *p* = 0.296; Figure 2A). This result indicated that the timing of motor execution was similar between each finger tapping. The averaged value of the participants’ answers regarding the ease of imagining tapping movements during MRI scanning was 2.46 (SD = 0.632) when collapsing four fingers.

**Figure 2:**
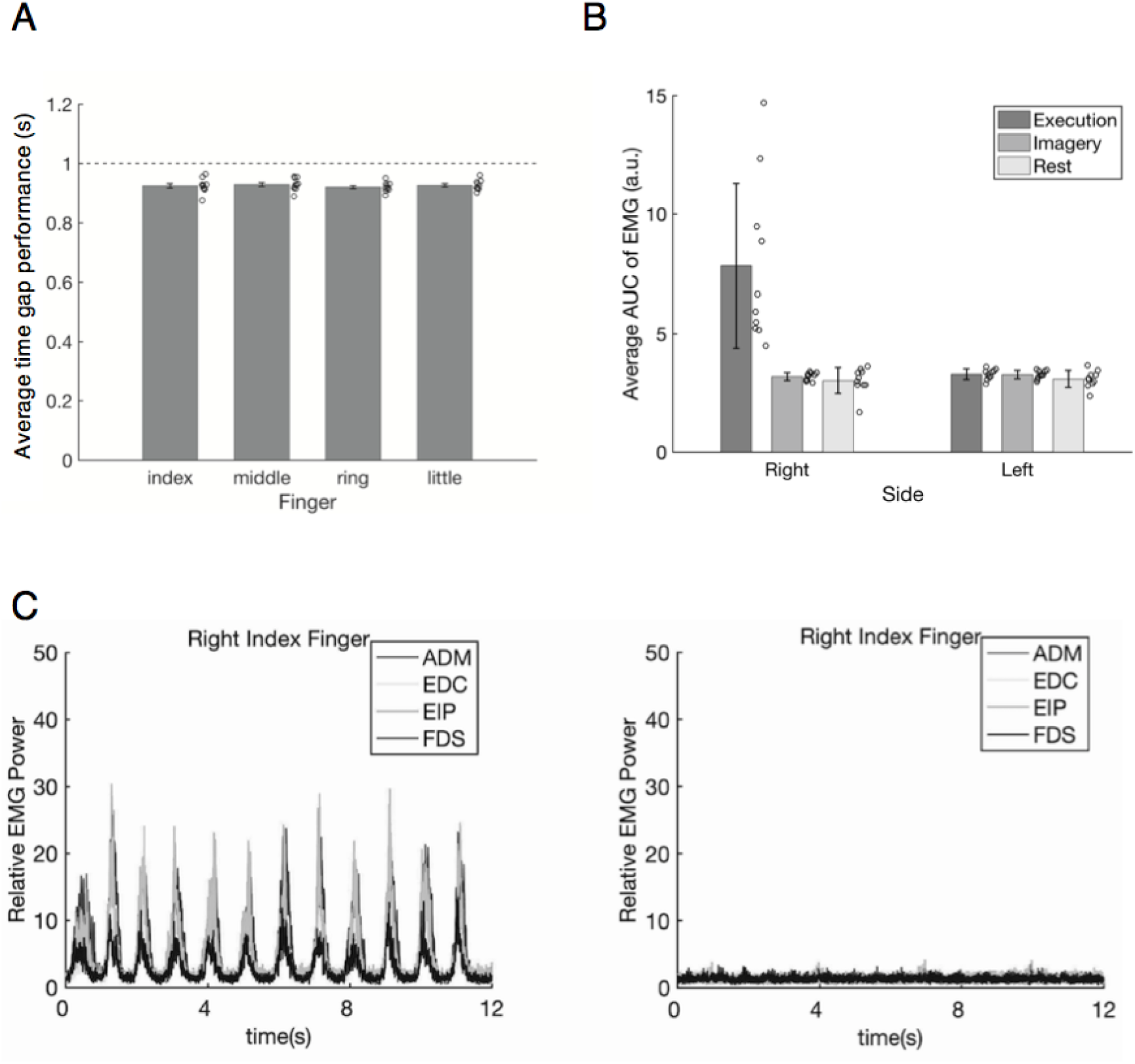
A) Time gap performance for each finger tapping. The horizontal dashed line indicates 1 s (1 Hz). Error bars indicate standard errors. Circles indicate individual data. B) The mean value of the area under the curve calculated from the electromyography (EMG) in each task for the respective side (left or right forearm). C) The typical EMG waveform extracted from the electrode attached to the right hypothenar (abductor digiti minimi: ADM), extensor digitorum communis (EDC), extensor indicis proprius (EIP), and flexor digitorum superficialis (FDS) muscles of one participant. The left graph shows the average waveform during index finger tapping calculated from the pooled execution task blocks, while the right graph shows that of the pooled imagery task blocks.

### EMG data analysis

For each combination of sides (left or right forearm) and sessions (execution, imagery, and fixation), we summed the area under the curve (AUC) of the EMG waveforms obtained from four muscles. The rest period of the fixation session corresponded to the average timing of the respective task blocks of all the execution and imagery task sessions. Because there were four execution task sessions and four imagery task sessions but only one fixation session, we averaged the AUC sum across the sessions for each task. We divided the original AUC values of the EMG by 10,000 to simplify calculations and processing (Figure 2B, C). See the supplementary material for details on the EMG recording and processing. The mean (SD) values of the execution, imagery, and rest blocks for the right arm were 7.828 (3.443), 3.186 (0.167), and 3.014 (0.550), respectively, and those for the left arm were 3.288 (0.220), 3.271 (0.176), and 3.082 (0.362), respectively.

Two-way within-participants ANOVA of the task condition (execution vs. imagery vs. rest) and side (left vs. right forearm) was performed. We detected a significant interaction (*F* (2, 18) = 17.759, *_p_*η*^2^* = 0.664, *p* < 0.001), and post hoc analysis revealed a significant simple main effect of side on the execution task condition (*F* (1, 27) = 50.761, *p* < 0.001) and a significant simple main effect of the task condition on the right side (*F* (2, 36) = 36.180, *p* < 0.001). Multiple comparison tests using Tukey’s honestly significant difference test indicated that for the right side, the AUC sum of the execution task was significantly larger than that of both the imagery task and rest period (Cohen’s *d*s > 1.359, *p*s < 0.01). Notably, there was no significant difference between the imagery task and the rest period (*d* = 0.325, *p* > 0.05). Thus, the results of MRI analysis regarding motor imagery cannot be regarded as epiphenomena of muscle activities.

### fMRI mass-univariate analysis

We calculated the average signal change rate of the respective combinations of ROIs and tasks (execution and imagery) with collapsing the four fingers (Figure 3A). These values were compared with 0 using a two-tailed one-sample *t*-test corrected using the Holm– Bonferroni procedure (Holm, 1979). In the execution task sessions, only the left hand-M1 (mean = 0.472, SD = 0.261) showed significant values higher than 0 (*t* (9) = 5.714, *d* = 1.905, *p* < 0.001; for the bilateral SMA (mean = 0.144, SD = 0.294) and visual areas (mean = 0.031, SD = 0.267), *t*s (9) < 1.545, *d*s < 0.515, *p*s > 0.157). In the imagery task sessions, the bilateral visual area (mean = −0.371, SD = 0.272) showed values significantly lower than 0 (*t* (9) = −4.324, *d* = 1.441, *p* < 0.002), while bilateral SMA (mean = 0.359, SD = 0.254) showed values significantly higher than 0 (*t* (9) = 4.478, *d* = 1.493, *p* < 0.002), but no significant values were shown for the left hand-M1 (mean = 0.036, SD = 0.289, *t* (9) = 0.394, *d* = 0.131, *p* = 0.703; see also Supplementary Figure 1 for results of mass-univariate group analyses). A consistent result was obtained when the rest block was modeled (see Supplementary Figure 2).

**Figure 3:**
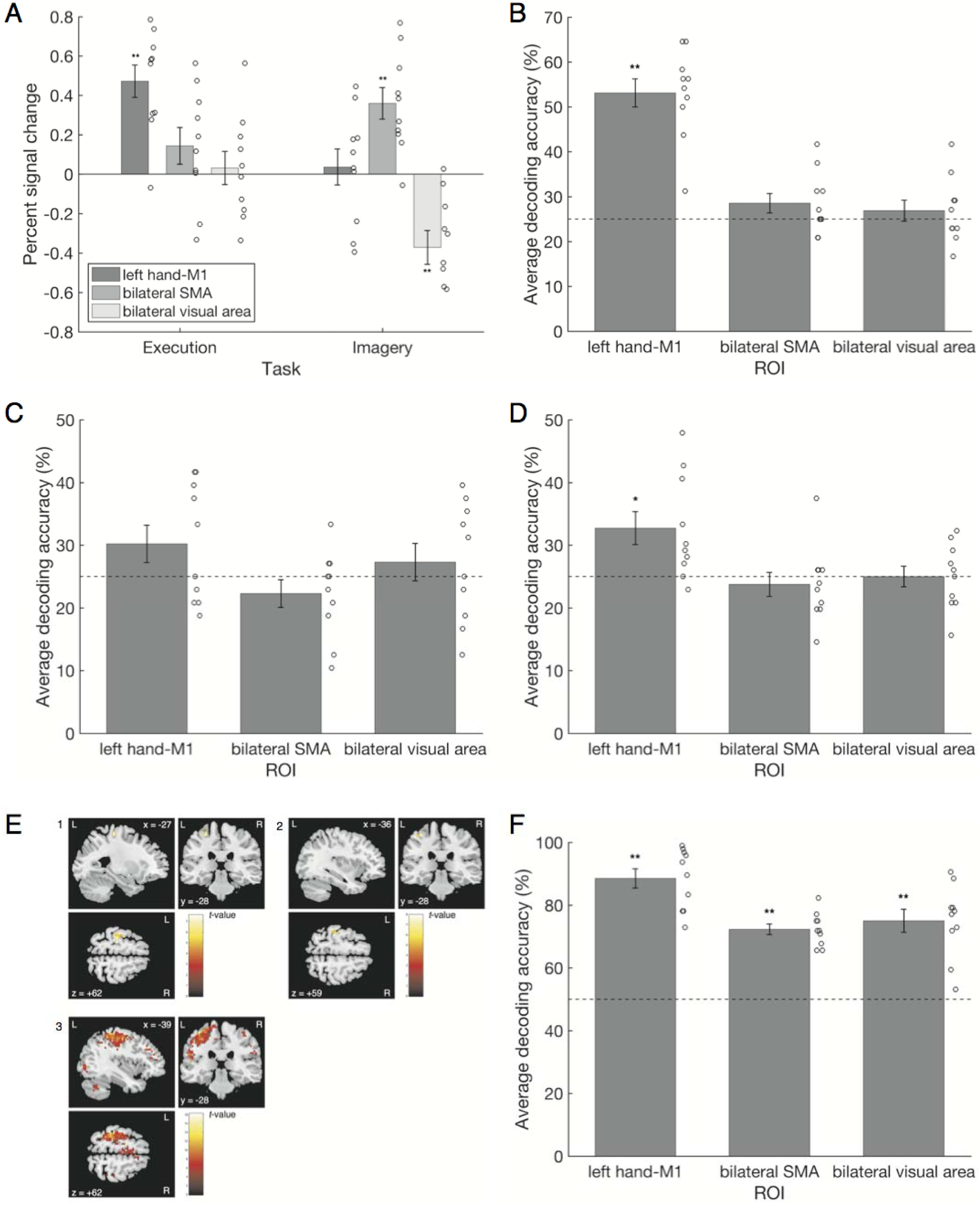
A) The average signal change rate of each ROI for each task session. B) The average decoding accuracy of each ROI in the within-execution classifications. C) Mean decoding accuracy of each ROI in the within-imagery classifications. D) Average decoding accuracy of each ROI calculated from two cross-classification directions. E) The results of the searchlight analyses for three types of classifications. The colored area is the detected cluster. L indicates the left hemisphere, while R indicates the right hemisphere. E-1) The result of the searchlight analysis for the within-execution classification with an uncorrected threshold of *p* < 0.0005, k ≥ 10. E-2) The result of the searchlight analysis for cross-classification between the execution and imagery task sessions with an uncorrected threshold of *p* < 0.0005, k ≥ 10. E-3) The result of the searchlight analysis for the classification of two modalities (execution and imagery) with an uncorrected threshold level of k ≥ 10, *p* < 0.0005. F) Average decoding accuracy of each ROI for the two modality classifications: execution and imagery. Error bars indicate standard errors. The horizontal dashed line indicates chance. Circles indicate individual data. *: *p* < 0.05; **: *p* < 0.01.

### Multi-voxel classification analysis

ROI analysis was divided into within-classifications and a cross-classification. Using the activation pattern of the left hand-M1 during motor execution (mean = 53.125, SD = 9.931), we were able to classify which finger was tapped more correctly than the chance-level of 25% (*t* (9) = 8.96, *d* = 2.985, *p* < 0.001; Figure 3B) from a within-execution classification. The classification accuracies for the bilateral SMA (mean = 28.542, SD = 6.878) and visual areas (mean = 26.875, SD = 7.379) were not significant (*t*s (9) < 1.63, *d*s < 0.543, *p*s > 0.138). In the within-imagery analysis, no ROI showed significantly higher classification accuracies than chance (means (SDs) of left hand-M1, bilateral SMA, and bilateral visual areas were 30.208 (9.433), 22.292 (6.948), and 27.292 (9.443), respectively; all ROIs were not significant: *t*s (9) < 1.75, *d*s < 0.582, *p*s > 0.115; Figure 3C). That is, we could correctly classify the type of motor execution using the activities of the left hand-M1 during actual movements, but we could not classify it in terms of motor imagery. The average classification accuracy of the two cross-classification directions, i.e., trained with the execution task sessions and tested with the imagery task sessions and vice versa, was significantly higher than the chance-level of 25% in the left hand-M1 (mean = 32.708, SD = 8.307, *t* (9) = 2.93, *d* = 0.978, *p* = 0.0166; Figure 3D). In other words, we could correctly classify the content of motor execution/imagery across tasks using the activities of the left hand-M1. When we performed statistical tests on these two directions separately, the direction where a classifier was trained with imagery data and tested with execution data showed a significantly higher than chance classification accuracy (mean (SD) = 35.833 (10.149), *t* (9) = 3.375, *p* = 0.008, *d* = 1.125), while the classification accuracy of the reverse direction was not significant (mean (SD) = 29.583 (8.437), *t* (9) = 1.718, *p* = 0.120, *d* = 0.573). The average classification accuracies of the bilateral SMA (mean = 23.75, SD = 6.05) and visual areas (mean = 25, SD = 5.243) did not differ significantly from chance (|*t*s (9)| < 0.655, *d*s < 0.218, *p*s > 0.530). The cerebral white matter and CSF also showed non-significant decoding accuracy (Supplementary Figure 3). The nonparametric permutation test had a more liberal *p*-value threshold than the parametric *t*-test. The results of the permutation tests supported the results of the parametric tests. The searchlight analysis was also divided into within-classifications and a cross-classification. For the within-execution classification, the left cluster consisting of BA4 and BA6 showed significantly higher-than-chance classification accuracies with an uncorrected threshold level of *p* < 0.0005, k ≥ 10 (Figure 3E-1; Table 1). We did not correct the threshold because the purpose of the searchlight analysis was not to find a significant location in the whole brain but to confirm the result of the ROI analysis. No cluster showed significantly higher-than-chance classification accuracy with the same threshold settings for within-imagery classification. The left cluster, consisting of BA4 and BA3, showed a significantly higher-than-chance classification accuracy at the same threshold settings for the combined two cross-classification directions (Figure 3E-2; Table 1).

**Table 1:**
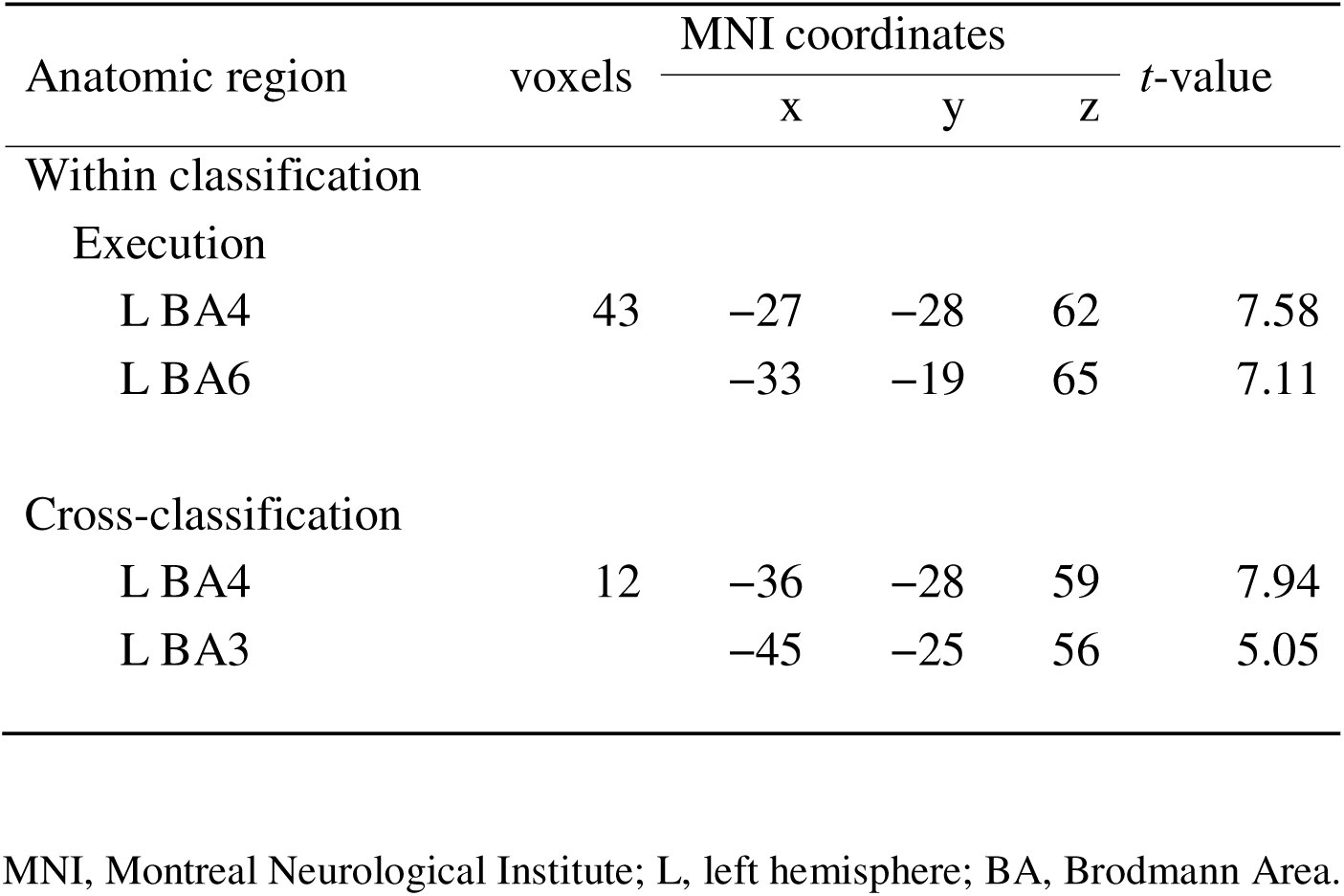
Anatomical regions, coordinates of peak voxel, and *t*-values of observed activations in the searchlight analysis.

| Anatomic region | voxels | MNI coordinates |  |  | *t*-value |
| --- | --- | --- | --- | --- | --- |
|  |  | x | y | z |  |
| Within classification |  |  |  |  |  |
| Execution |  |  |  |  |  |
| L BA4 | 43 | −27 | −28 | 62 | 7.58 |
| L BA6 |  | −33 | −19 | 65 | 7.11 |
| Cross-classification |  |  |  |  |  |
| L BA4 | 12 | −36 | −28 | 59 | 7.94 |
| L BA3 |  | −45 | −25 | 56 | 5.05 |

MNI, Montreal Neurological Institute; L, left hemisphere; BA, Brodmann Area.

Next, we attempted between-modality classification. For these analyses, we collapsed the data from the four fingers and tried to determine whether they came from the execution task session or the imagery task session. Chance level was 50%. The ROI analysis showed that the decoding accuracies were significantly higher than chance for all ROIs (means (SDs) of left hand-M1, bilateral SMA, and bilateral visual areas were 88.542 (9.635), 72.292 (5.271), and 75 (11.61), respectively; *t*s (9) > 3.98, *d*s > 1.33, *p*s < 0.004; Figure 3F). In the searchlight analysis, the left cluster consisting of BA4 and BA6 showed classification accuracies that were significantly higher than chance, with an uncorrected threshold level of k ≥ 10, *p* < 0.0005. Its peak voxel belonged to the left M1 (Figure 3E-3).

### Representational similarity analysis

RSA was performed on the left hand-M1 because it was the only ROI that showed significant cross-classification accuracy. For each participant, the RDM was estimated as an 8 by 8 (four executed and imagined tappings) matrix of dissimilarity. The average RDM (Figure 4A) was calculated from the RDMs of all the participants in the execution and imagery tasks. One of our predictions focused on whether motor imagery in the RDM would have characteristics similar to those found by Ejaz et al. (2015) and Ogawa et al. (2019) in previous motor execution studies. Ogawa et al. (2019) found that two adjacent fingers had a smaller distance than two more distant fingers, i.e., two fingers with the other finger in between, such as the index and ring finger pair. Ejaz et al. (2015) reported that the middle-ring finger pair or ring-little finger pair had a smaller distance than the other finger pairs.

**Figure 4:**
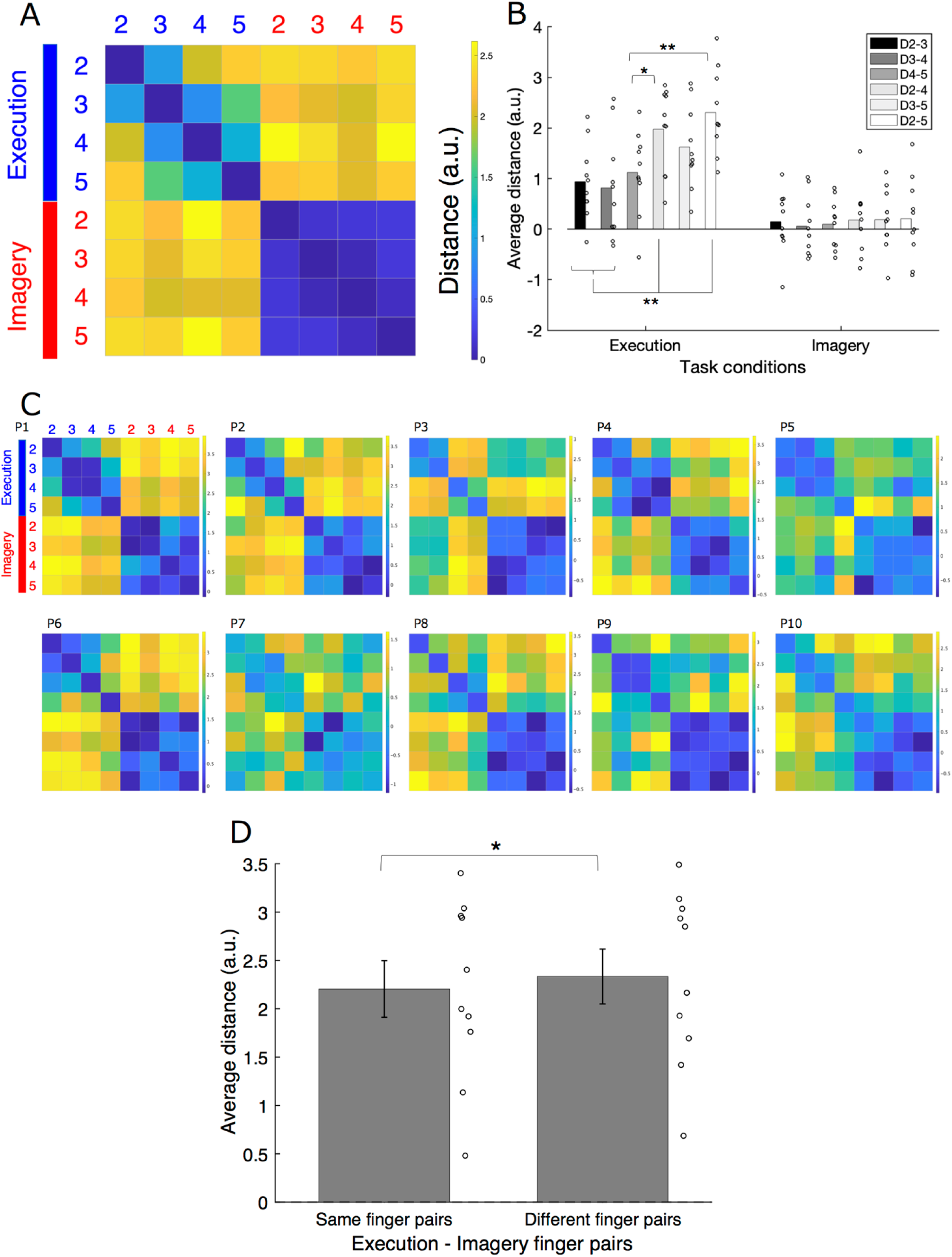
A) The RDM averaged over all participants. Each cell represents the dissimilarity between the activity patterns of two fingers as a distance. 2, the index finger; 3, middle finger; 4, ring finger; and 5, little finger. Blue text, execution; red text, imagery. B) Average distances of finger pairs within the task. C) RDM of each participant. Within-execution finger pairs exhibited a relatively stable pattern across participants, with the distances between adjacent finger pairs being smaller than those between distant finger pairs. In contrast, the patterns of the within-imagery finger pairs were less stable across participants. D) Average distances of diagonal (same finger) pairs and nondiagonal (different finger) pairs in the motor execution and imagery combination domain in the average RDM. Circles indicate individual data. Error bars show standard errors. *: *p* < 0.05, **; *p* < 0.01.

A two-way within ANOVA was performed on the distance of the six within-task finger pairs for both the execution and imagery tasks (two tasks × six finger pairs; Figure 4B; see also 4C for individual RDMs). Means (SDs) of index-middle finger pair, middle-ring finger pair, ring-little finger pair, index-ring finger pair, middle-little finger pair, and index-little finger pair in the execution task were 0.937 (0.745), 0.814 (1.037), 1.119 (0.823), 1.977 (0.825), 1.621 (0.816), and 2.305 (0.824), respectively, while the means (SDs) of the same pairs in the imagery task were 0.146 (0.618), 0.054 (0.592), 0.094 (0.488), 0.178 (0.669), 0.187 (0.614), and 0.204 (0.811), respectively. If the execution and imagery task share a common feature in representational geometry, there should be main effects of finger pair, with no interaction. In the results, both the main effects of task and finger pair were significant (*F* (1, 9) = 37.764, *p* < 0.001, *_p_*η*^2^* = 0.808, *F* (5, 45) = 5.975, *p* < 0.001, *_p_*η*^2^* = 0.399, in order). The interaction was also significant (*F* (5, 45) = 3.64, *p* = 0.008, *_p_*η*^2^* = 0.288), and post hoc analysis showed the significant simple main effects of the task on each finger pair (*F*s (1, 54) > 4.97, *p*s < 0.03); for all finger pairs, the execution task showed greater distance than the imagery task. A significant simple main effect of finger pair on the execution task was also observed (*F* (5, 90) = 9.333, *p* < 0.001), whereas the simple main effect of finger pair on the imagery task was not significant (*F* (5, 90) = 0.088, *p* = 0.994). Multiple comparison tests using Tukey’s honestly significant difference showed that each pair of adjacent fingers had a smaller distance than the index-ring finger pair and the index-little finger pair in the execution task (*d*s > 0.69, *p*s < 0.05). These results showed that the dissimilarity of finger pairs in the execution task was generally higher than that in the imagery task. Furthermore, as Ogawa et al. (2019) indicated, the dissimilarity in the neural representations of adjacent fingers was lower than that of distant fingers in the representational geometry of the execution task. In contrast, this feature was not statistically supported by the representational geometry of the imagery task.

In addition, we tested the other prediction regarding the representational geometry of motor execution and imagery. For each participant, we averaged the cross-validated Mahalanobis distance between the neural representation of the executed finger movement and that of the imagined same-finger movement from the index finger to the little finger. We also averaged the distances of all other executed-imagined finger-movement pairs. In other words, we separately calculated the mean distance value of the diagonal and nondiagonal pairs in the region of the execution-imagery combination in the RDM. The two means were compared using a two-way paired *t*-test. The results showed that the mean distance (SD) of the same finger pairs (2.204 (0.926)) was significantly shorter than that of different finger pairs (2.334 (0.897)) (*t* (9) = −2.38, *d* = 0.80, *p* = 0.040) (Figure 4D).

### Replication study

Using a two-way *t*-test, we compared the average values of the participants’ responses regarding the ease of imagery during the imagery session between the original and replication studies and found no significant differences (first study, mean = 2.46, SD = 0.632; replication study, mean = 2.174, SD = 0.606; *t* (22) = 1.120, *d* = 0.486, *p* = 0.274).

Consistent with the MVPA results in the first study, the mean classification accuracy of the two cross-classification directions was significantly higher than chance (25%) in the left hand-M1 (mean = 37.5, SD = 7.182, *t* (13) = 6.51, *d* = 1.810, *p* < 0.0001; for the bilateral SMA and visual areas, means (SDs) were 25.744 (3.355) and 27.083 (4.971), respectively, *t*s (13) < 1.57, *d*s < 0.430, *p*s > 0.141; Figure 5A). Similar to the first study, the results of the permutation tests supported these findings. We correctly classified the content of motor execution/imagery across tasks using left hand-M1 activity. Each direction of cross-classification achieved a significantly above chance accuracy (means (SDs) = 39.286 (8.796) [trained with motor imagery and tested with motor execution] and 35.714 (6.216) [reverse direction], *t*s (13) = 6.077, 6.45, *p*s < 0.001, *d*s > 1.686).

**Figure 5:**
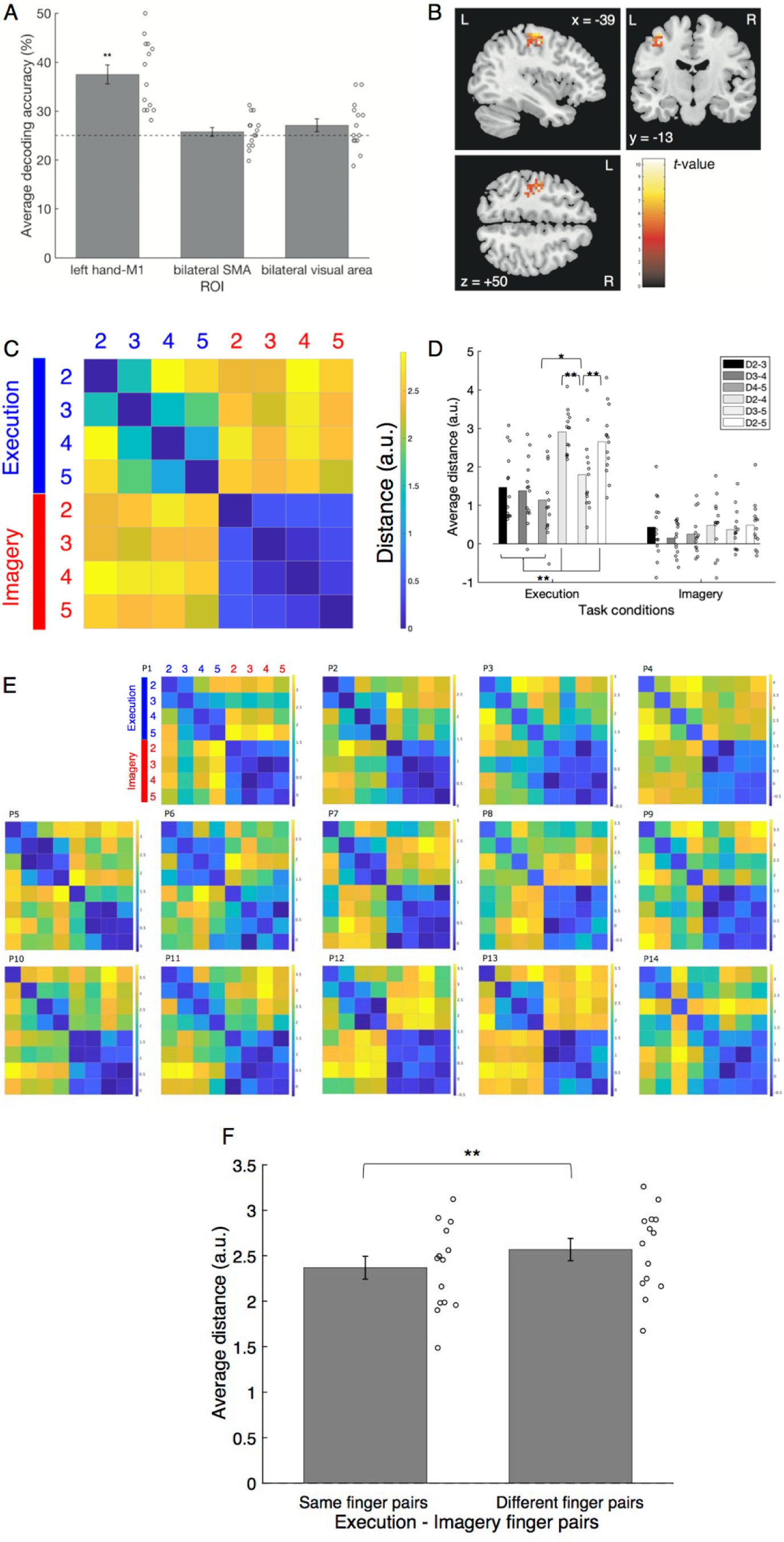
A) The average decoding accuracy of each ROI calculated from two cross-classification directions obtained in the replication study. The horizontal dashed line indicates chance-level. B) The result of the searchlight analysis for cross-classification between the execution and imagery in the replication study with an uncorrected threshold level of *k* ≥ 10 (*p* < 0.0005). L indicates the left hemisphere, and R indicates the right hemisphere. C) The RDM averaged across participants, in which each cell shows the distance (dissimilarity) between the activity patterns of two fingers. 2, the index finger; 3, middle finger; 4, ring finger; and 5, little finger. D) Mean distances of within-task finger pairs in each task. E) Individual RDMs of each participant. F) Average distances of diagonal (same finger) pairs and nondiagonal (different finger) pairs in the motor execution and imagery combination domain in the average RDM. Circles indicate individual data. Error bars show standard errors. *: *p* < 0.05; **: *p* < 0.01.

We also conducted a cross-classification searchlight analysis. The left cluster located on BA4 showed a significantly higher than chance classification accuracy with the same threshold settings as in the first study (Figure 5B; Table 2).

**Table 2:**
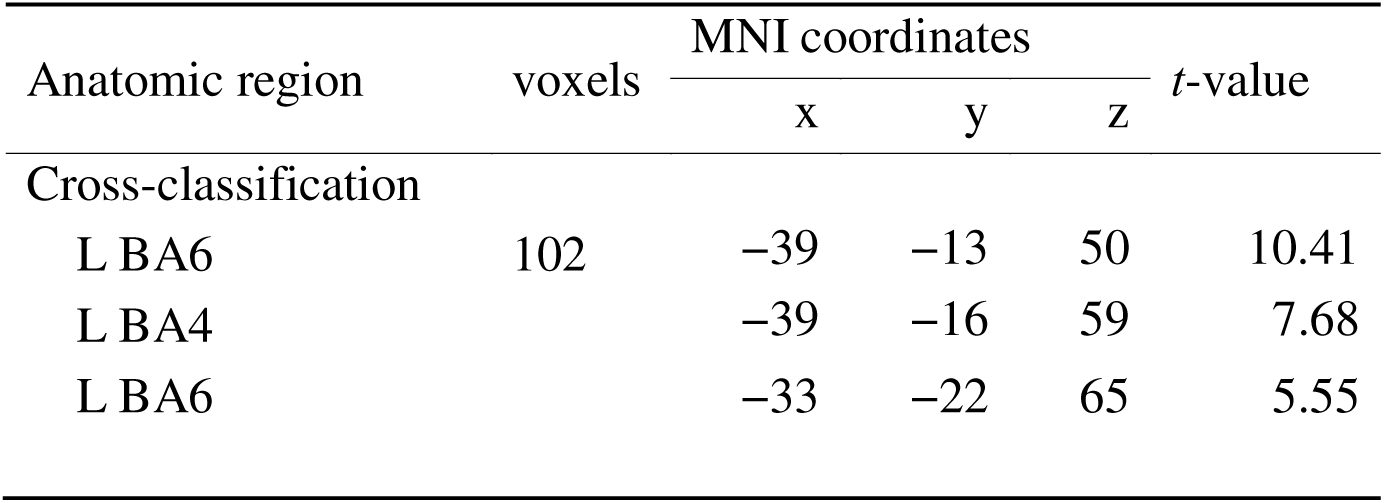
Anatomical regions, coordinates of peak voxel, and *t*-values of observed activations in the searchlight analysis of the replication study. MNI, Montreal Neurological Institute; L, left hemisphere; BA, Brodmann Area.

| Anatomic region | voxels | MNI coordinates |  |  | *t*-value |
| --- | --- | --- | --- | --- | --- |
|  |  | x | y | z |  |
| Cross-classification |  |  |  |  |  |
| L BA6 | 102 | −39 | −13 | 50 | 10.41 |
| L BA4 |  | −39 | −16 | 59 | 7.68 |
| L BA6 |  | −33 | −22 | 65 | 5.55 |

RSA targeting the left hand-M1 was also performed, and the RDM was averaged across participants in a manner similar to that in the first study (Figure 5C). Means (SDs) of the index-middle finger, middle-ring finger, ring-little finger, index-ring finger, middle-little finger, and index-little finger pairs in the execution task were 1.47 (0.878), 1.381 (0.83), 1.139 (0.958), 2.908 (0.557), 1.797 (0.962), and 2.652 (0.867), respectively, while the means (SDs) of the same pairs in the imagery task were 0.435 (0.764), 0.15 (0.429), 0.255 (0.516), 0.481 (0.716), 0.369 (0.52), and 0.486 (0.61), respectively. A two-way within ANOVA (two tasks × six finger pairs) was performed on the distance of the finger pairs (Figure 5D, see also 5E as individual RDMs). The results were similar to those of the first study. Both the main effects of task and finger pair were significant (*F* (1, 13) = 45.94, *p* < 0.001, *_p_*η*^2^* = 0.779, *F* (5, 65) = 14.359, *p* < 0.001, *_p_*η*^2^* = 0.525, respectively). The interaction was also significant (*F* (5, 65) = 8.561, *p* < 0.001, *_p_*η*^2^* = 0.397), and post hoc analysis showed the significant simple main effects of the task on each finger pair (*F*s (1, 78) > 8.75, *p*s < 0.04), with all the execution finger pairs having a greater distance than the imagery finger pairs. A significant simple main effect of the finger pair on the execution task was also observed (*F* (5, 130) = 22.277, *p* < 0.001), whereas the simple main effect of the finger pair on the imagery task was not significant (*F* (5, 130) = 0.256, *p* = 0.569). Multiple comparison tests using Tukey’s honestly significant difference showed that each pair of adjacent fingers had a smaller distance than the index-ring finger pair and the index-little finger pair in the execution task. Furthermore, the ring-little finger pair had a smaller distance than the middle-little finger pair, while the middle-little finger pair had a smaller distance than the index-ring finger pair and the index-little finger pair (*d*s > 0.668, *p*s < .05). These results were not contradictory to those of the first study or the effect of adjacent fingers, as suggested by Ogawa et al. (2019). Additionally, these results suggest that the ring-little finger pair has a particularly low dissimilarity among finger pairs, for example, partially similar to that suggested by Ejaz et al. (2015). In addition, as in the first study, the respective mean distances for the same (diagonal) finger pairs and different (nondiagonal) finger pairs in the region of the execution-imagery combination in the RDM were calculated for each participant. A two-way paired *t*-test showed that the mean distance (SD) of the same finger pairs (2.368 (0.469)) was significantly shorter than that of the different finger pairs (2.568 (0.457)) (*t* (13) = −3.45, *d* = 0.96, *p* = 0.004) (Figure 5F).

To visualize how finger tapping for motor execution and imagery are represented as brain activity patterns in the left hand-M1, we conducted a principal component analysis using classical multi-dimensional scaling (MDS) with the averaged RDM (n = 24), combining both the first and replicated studies (Figure 6). This figure clearly illustrates the relationship between execution and imagery in representational geometry by showing that the same fingers are closely located irrespective of modality.

**Figure 6:**
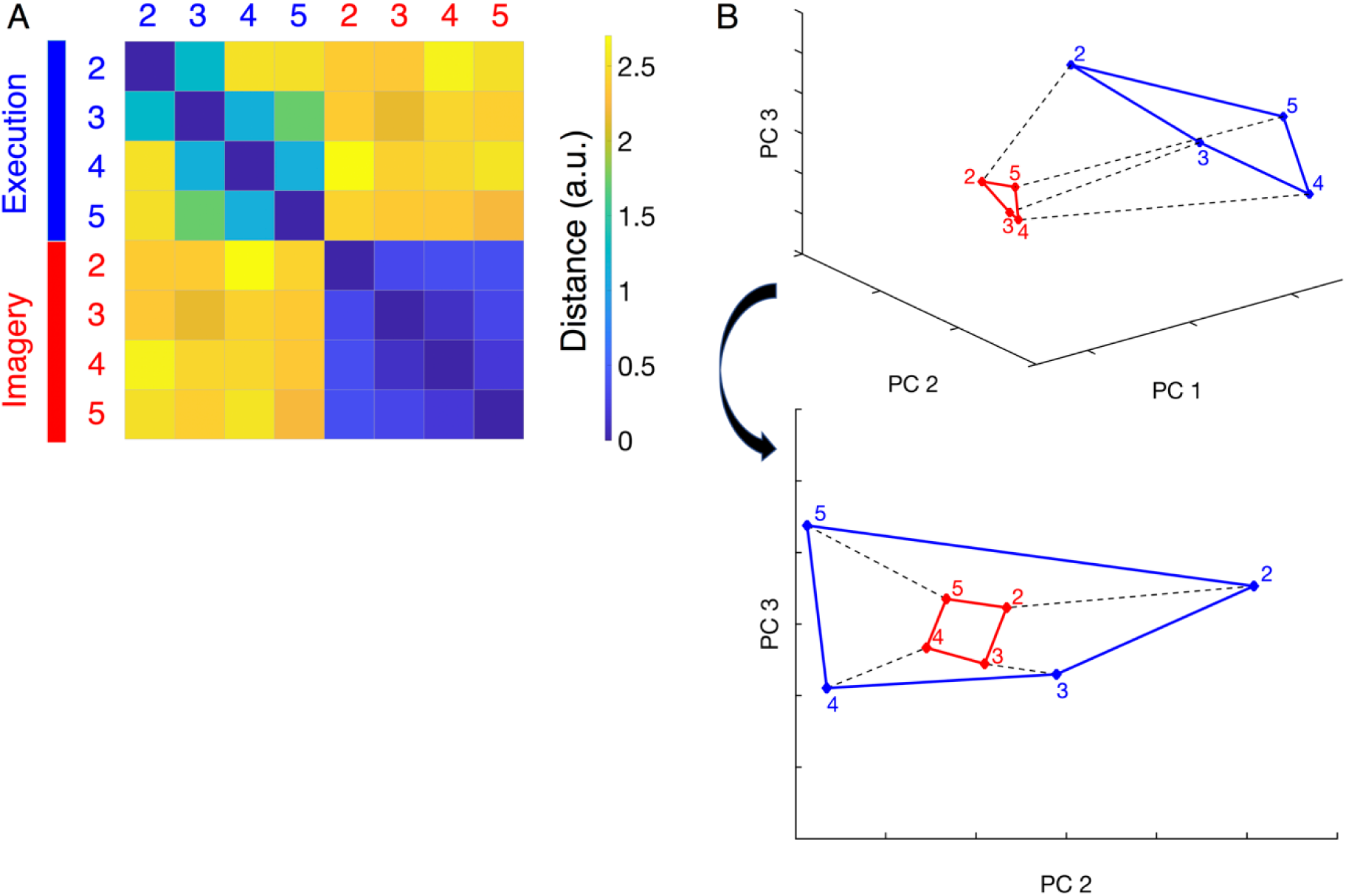
The representational geometry of the left hand-M1. A) The RDM averaged across 24 participants was drawn from each pair of executed and imagined finger movements. Each cell indicates the extent to which the activity patterns for two finger movements differ from each other. 2, the index finger; 3, middle finger; 4, ring finger; and 5, little finger. B) Plots of eight finger movements, i.e. four execution and four imagery, drawn by classical MDS. From PC1 to PC3, the eigenvalue was high in that order.

## Discussion

This study investigated whether the M1 relates to motor imagery similar in a manner similar to that of motor execution. The first study showed a significantly higher-than-chance accuracy with cross-classification between execution and imagery in the left hand-M1. The replication study successfully reproduced this result. Thus, we demonstrated that the contralateral hand M1 contains a common neural representation between motor execution and kinesthetic motor imagery.

Mass-univariate analysis of the first study revealed that the average activity of the left hand-M1 significantly increased during motor execution compared with the baseline activity at rest. In the MVPA, both the ROI and searchlight analyses detected the left M1 as above-chance-level classification accuracy for within-execution. In contrast, parametric *t*-tests in the within-imagery, searchlight, and mass-univariate analyses of the imagery task showed no significant results for the left M1. Since the M1 is associated with motor control of contralateral body parts (Heming et al., 2019), these results suggest that, in addition to the disparity in the overall amount of brain activity between motor execution and motor imagery, distinct patterns of brain activity emerge during motor execution of finger tappings that may not be as evident during motor imagery. In both the first and replication studies, RSA showed that the distance for each pair of fingers for motor execution was larger than that for motor imagery. These results support the interpretation that fingers can be clearly separated in the brain activity patterns of motor execution, whereas this separation may not be as clear in motor imagery.

To our knowledge, this is the first study to report successful cross-classification between motor execution and kinesthetic imagery in the contralateral hand M1. The cross-classification accuracy in a previous study (Zabicki et al., 2017; see also Monaco et al., 2020) remained at chance level for the M1. This difference may be due to methodological differences. Unlike the current study, Zabicki et al. did not use a functional localizer to define M1 as an ROI. Another factor is the hand movements used in previous studies: pointing to a target, squeezing, and extension-flexion. These actions differ in many aspects, including spatiality, timing, the presence of an action target, and effectors (Pilgramm et al., 2016). Pilgramm et al. (2016) recommended controlling the parameter of each action property to determine the type of information represented in each brain area. The same claim was made by Monaco et al. (2020), in which participants imagined grasping movements with a large or small object and reaching movements. The movements in our tasks were similar in terms of spatiality, action target, and timing, as shown in the timing performance in Figure 2A. In contrast, these actions differed in the motor effector, i.e.,which finger tap was executed/imagined. The presence of a target may also affect the content of the imagery. It is possible that the participants in the previous study imagined not only body parts but also the target object involved in the actions, whereas our participants generated more motor effector-focused imagery. If so, M1, which has a neural representation of the fingers (e.g., Ejaz et al., 2015; Ogawa et al., 2019), may have been more sensitive to the content of motor imagery in the present study.

Notably, the mass-univariate analysis revealed no increase in left hand-M1 activation during motor imagery. Therefore, even if the contralateral hand M1 showed no increased activation, it could still represent a finger of the imagined movement, which was partially similar to motor execution, especially in tasks where the imagined movements were of the same type and differentiated by the motor effector.

RSA clarified the representational geometric relationship between motor execution and imagery. Consistent with a previous study (Ogawa et al., 2019), the dissimilarity between adjacent fingers was lower than that between the other two distant fingers in both the first and replication studies, and the ring-little finger pair had particularly low dissimilarity in the replication study (Ejaz et al., 2015). However, this effect was not observed for the imagery task in either study (Figure 4B and 5D). The imagery task consistently produced more negative dissimilarity scores than did the execution task (small circles in Figure 4B and 5D). Negative scores mean that the pattern of activity for each finger is inconsistent, and thus left hand-M1 does not reliably encode differences between them (e.g., Boccia et al., 2021). Therefore, it appears that within the participants, the activity pattern of each imagined finger movement was less stable than that of the executed finger movement. One possible reason for this is that imagined finger movements lack tactile feedback from button presses. Previous studies have suggested that the M1 processes tactile sensations (e.g., see the case of fingers, Schroeder et al., 2018). Such sensory processing could also increase the differences in neural activity patterns between different fingers during the execution task.

We also analyzed the dissimilarity distance of two fingers between motor execution and motor imagery. In both the first and replication studies, the dissimilarity in brain activity patterns between motor execution and imagery of the same fingers was smaller than that between different fingers (Figures 4D and 5F). This relationship is clearly visualized by classical MDS based on the average RDM (Figure 6). Because motor execution and imagery were clearly separated along with PC1 (Figure 6B top view), the PC1 axis appeared to indicate the difference between motor execution and imagery. Ariani et al. (2022) reported that the motor planning for simple finger tapping has a representational geometry similar to its execution version. In the present study, the same fingers were the closest between motor execution and imagery (Figure 4D, 5F), although the effect of adjacent fingers was found only for motor execution (Figure 4C, 5D). Therefore, it could be said that the representational geometry of kinesthetically imagined finger tapping also has, in part, a “scaled-down version of the patterns during execution” (Ariani et al., 2022).

There are some possible reasons why the classification accuracy remained at the chance level in the within-imagery classification. First, the cross-classification used data from four task sessions of either execution or imagery, whereas the within-classification used only three task sessions as training data for each fold. Therefore, the classifier would have been better trained in the cross-classification than in the within-imagery classification. Second, as shown in Figure 6, the neural activity patterns of motor imagery may be closer and thus less distinguishable than those of motor execution. If so, the decoding accuracy should be higher for classifications trained with motor imagery and tested with a more discriminable motor execution than for the reverse direction. In the first study, the decoding accuracy was 35.833% and 29.583% for the former and latter classifications, respectively. The same pattern was observed in the replication study.

The MVPA showed chance-level decoding accuracy for the bilateral SMA and visual area (BA17 and BA18, respectively). Monaco et al. (2020) reported that within-imagery classification successfully decoded imagined movements in the early visual cortex. This difference could be attributed to the fact that they used visually guided movements, whereas we did not. The results of mass-univariate analysis showed increased activation only for the bilateral SMA during motor imagery, consistent with previous studies in which positive M1 activation was rarely detected and the relationship between the SMA and motor imagery was consistently identified (Hanakawa, 2016; Hétu et al., 2013). Kasess et al. (2008) suggested that the SMA suppresses M1 activation during motor imagery. Our results were consistent with this suggestion. The bilateral visual area showed negative activation during motor imagery compared to implicit baseline and resting-state activation (Supplementary Figure 2). Amedi et al. (2005) discovered that visual imagery suppresses the auditory cortex. This finding is consistent with the negative activation of the visual cortex observed in the present study. Further studies are required to clarify this phenomenon.

The present study had some limitations. First, it is possible that our main results in M1 were not due to the finger representation itself but rather to the spatial information of the target. Fabbri et al. (2014) reported that the M1 may represent the target direction of proprioceptively guided reaching movements via MVPA. However, the way the participants grasped the target had a greater effect on M1 than reach direction. Therefore, the likelihood that our results reflect only the difference in target spatial position without any motor component is relatively low. Second, it was not easy to prove that the participants generated kinesthetic motor imagery. Hanakawa et al. (2003) developed a task in which participants could not answer a question without using imagery. However, their task required multi-finger tapping and was therefore incongruent with the purpose of the present study. Our participants first performed the execution sessions, followed by the imagery sessions. We expected this order to familiarize the participants with the kinesthetic sensation of finger tapping and make it easier to generate kinesthetic motor imagery.

Recently, Persichetti et al. (2020) suggested that M1 was related to motor imagery using vascular space occupancy. They found that the superficial layers of M1, which received cortical input from other regions related to motor planning, showed an increase in cerebral blood volume related to imagery. In contrast, deep layers, which sent output to the spinal cord for motor execution, did not show such an increase. They did not directly examine the spatial pattern of cortical activation between executed and imagined movements. Although our study did not distinguish between the superficial and deep layers of the M1, based on the claim by Persichetti et al. (2020), our findings are most likely attributable to the superficial M1 layers. It is also important to note that the activation of M1 corticospinal neurons, which are mainly located in the M1 deep layers (e.g., in mice, Oswald et al., 2013), generally results in the release of motor signals to the spinal cord and execution of movement. Therefore, it is likely that the superficial layers of M1, rather than the deeper layers, represent imagined movements similar to executed movements.

In conclusion, we found that motor imagery has a spatial neural representation similar to execution in the contralateral hand M1, at least for the same finger. The representational geometry of finger movements during motor imagery may be partially a “scaled-down version of the patterns during execution” (Ariani et al., 2022). Future studies should investigate whether motor execution and imagery have similar representational geometries for different two-finger combinations, such as the effects of adjacent fingers (Ejaz et al., 2015; Ogawa et al., 2019).

## Acknowledgments

We thank Dr. Takashi Nagamine for helpful comments on an early version of this manuscript and Yusuke Haruki for helping with the experiment. We also thank Editage (www.editage.jp) for English language editing. The manuscript was written with the grammatical support of the AI-powered writing assistant, DeepL Write.

## Data and code availability statements

All the data are available upon reasonable request from the corresponding author.

## Supplementary materials

### EMG recording and processing

The electromyography (EMG) was measured by BrainAmp ExG MR (BRAIN PRODUCTS) and recorded with the Recorder (BRAIN PRODUCTS) with a sampling rate of 5000 Hz, low cutoff and DC, high cutoff of 1000 Hz. During EMG recording, the EMG clock was synchronized with the clock of an MRI scanner. Offline processing (e.g., noise filtering) of the EMG data was done using Analyzer2 (BRAIN PRODUCTS) in the following order: 1) MR noises were excluded, 2) EMG data were processed using the IIR (zero phase-shift Butterworth) filter with low cutoff: 5 Hz; high cutoff: 100 Hz; Notch: 50 Hz; Bandreject: 17, 33, 50, 67 Hz; bandwidth: 4 Hz, 3) EMG data were downsampled from 5000 Hz to 500 Hz, and 4) EMG data were rectified.

As written in the EMG analysis of results, the analyzed EMG data were extracted from the task blocks of the execution and imagery sessions, and the corresponding timing of the fixation session as a rest period. Generally, some participants can relax well when task onset comes, but others cannot. Cared about such an individual difference, each EMG data extracted from the target block was standardized by the standard deviation of the EMG values that were obtained from the 300 ms period just before each target block.

### Result of the functional localizer

**Supplementary Table 1:**
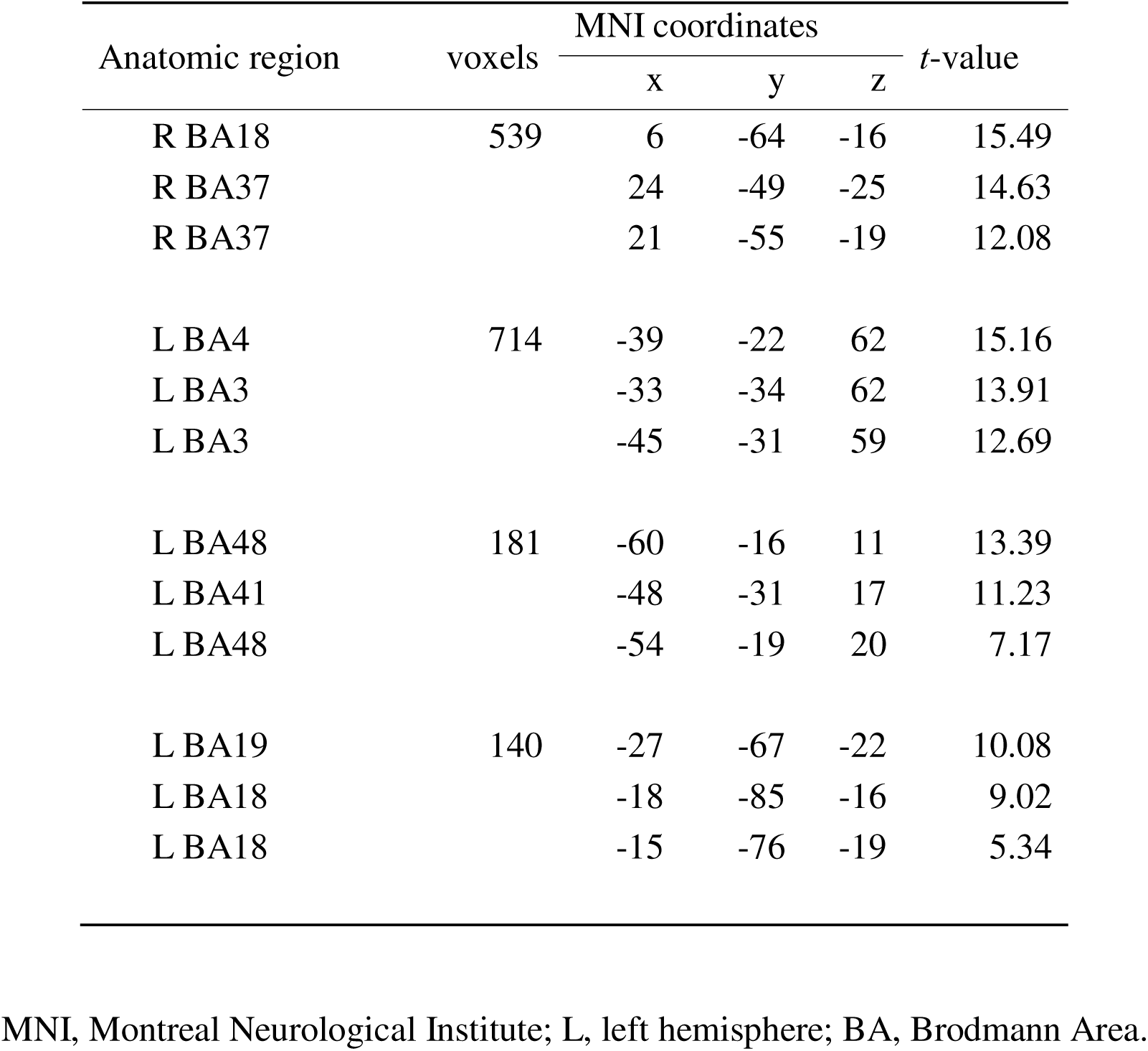
Anatomical regions, coordinates of peak voxel, and *t*-values of observed activations in the functional localizer of the first study.

### Results of the mass-univariate group analysis (Figure caption)

**Supplementary Figure 1:**
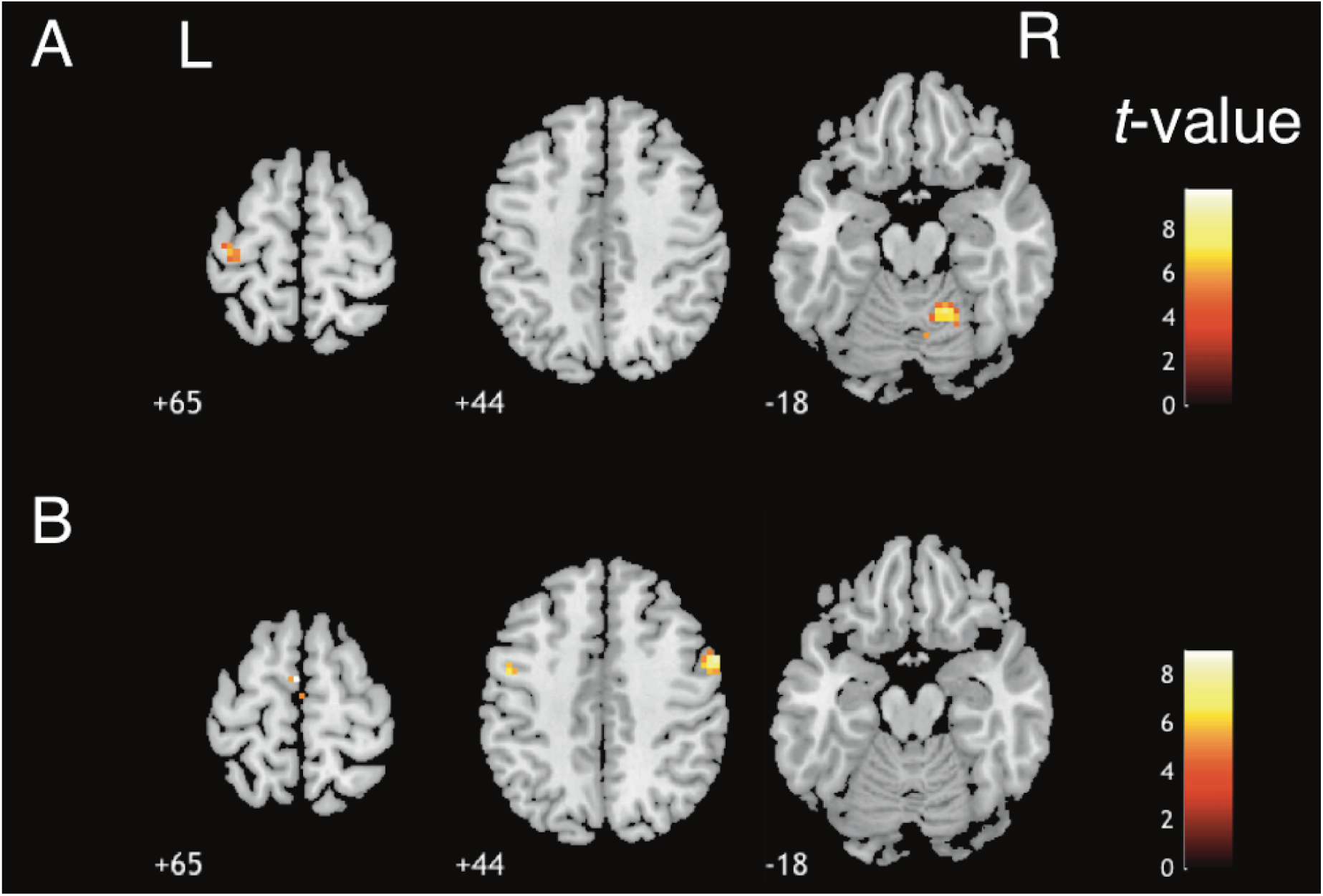
(A) The results of mass-univariate group analysis for the execution task session of the first study, with an uncorrected threshold level of *k* ≥ 10 (*p* < 0.0005). The detected left cluster, consisting of BA4 (i.e., X = −39, Y = −19, Z = +65, *t* = 4.862) and BA6, is indicated by the colored area. The detection of BA4 was to be expected, because this region relates to motor control of the contralateral body parts (e.g., Heming et al., 2019). (B) The results of mass-univariate group analysis for the imagery task session in the original study with an uncorrected threshold level of *k* ≥ 10 (*p* < 0.0005). The detected left cluster included BA6 (i.e., X = −3, Y = −4, Z = +65, *t* = 8.981) and is indicated by the colored area, but no voxels from BA4 were detected. SMA is frequently reported as an active region during motor imagery (e.g., Hanakawa, 2016), while M1 activation is rarely reported (Hétu et al., 2013). These results correspond to those in previous studies.

**Supplementary Figure 2:**
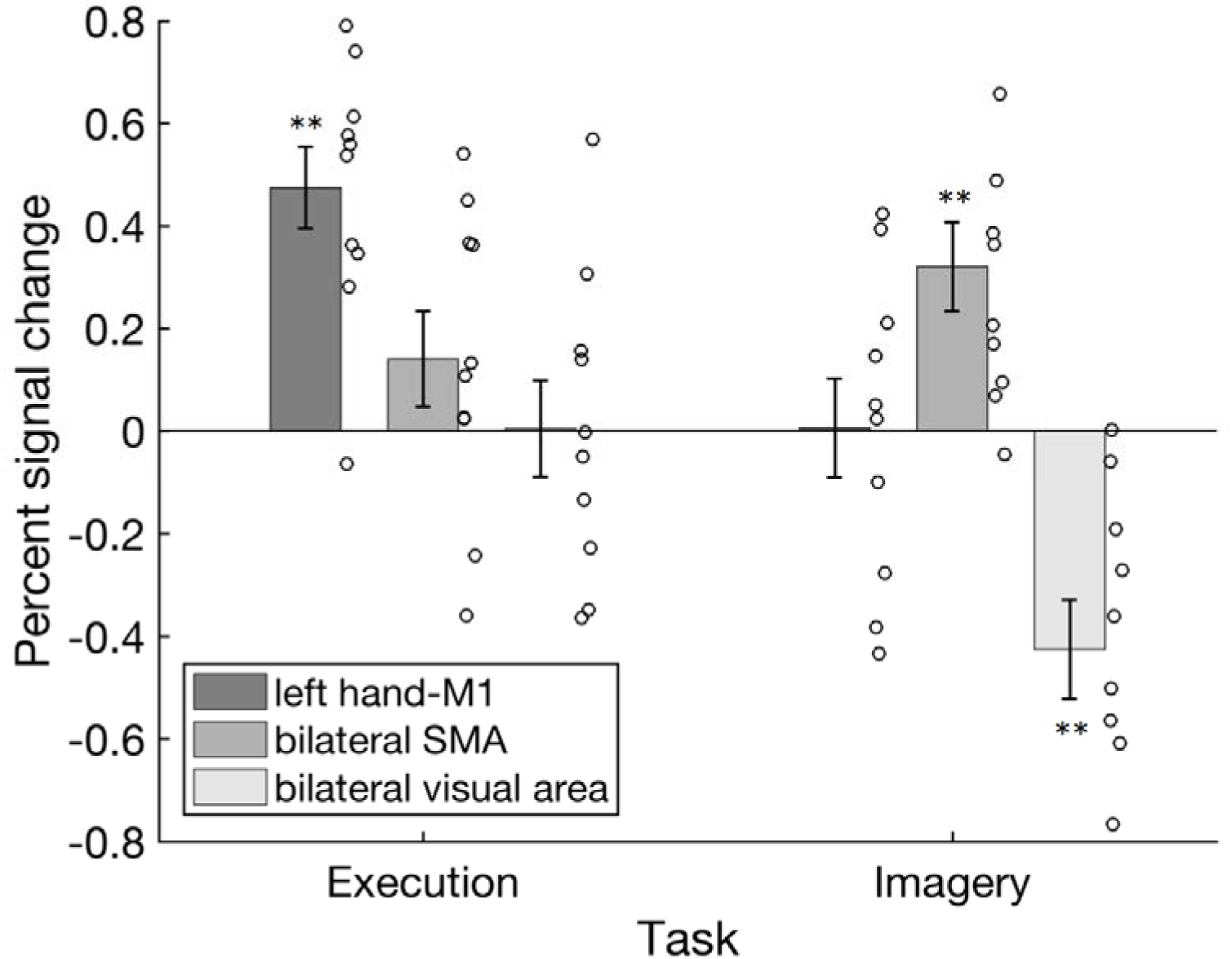
The result of the mass-univariate analysis with modeling of rest blocks: The average signal-change rate of each ROI in each task session. Error bars show standard errors. Circles indicate single data. **: *p* < 0.01. Consistent with the result of the mass-univariate analysis without modeling rest, significant differences from rest were found at three locations. Left hand-M1 in motor execution task (*t* (9) = 5.97, *p* < 0.01 (two-tailed), *d* = 1.99), bilateral SMA (*t* (9) = 3.7, *p* < 0.01, *d* = 1.23), and bilateral visual area (*t* (9) = −4.43, *p* < 0.01, *d* = 1.48) in motor imagery task. These results supported our conclusion based on the result of the mass-univariate analysis with modeling rest.

**Supplementary Figure 3:**
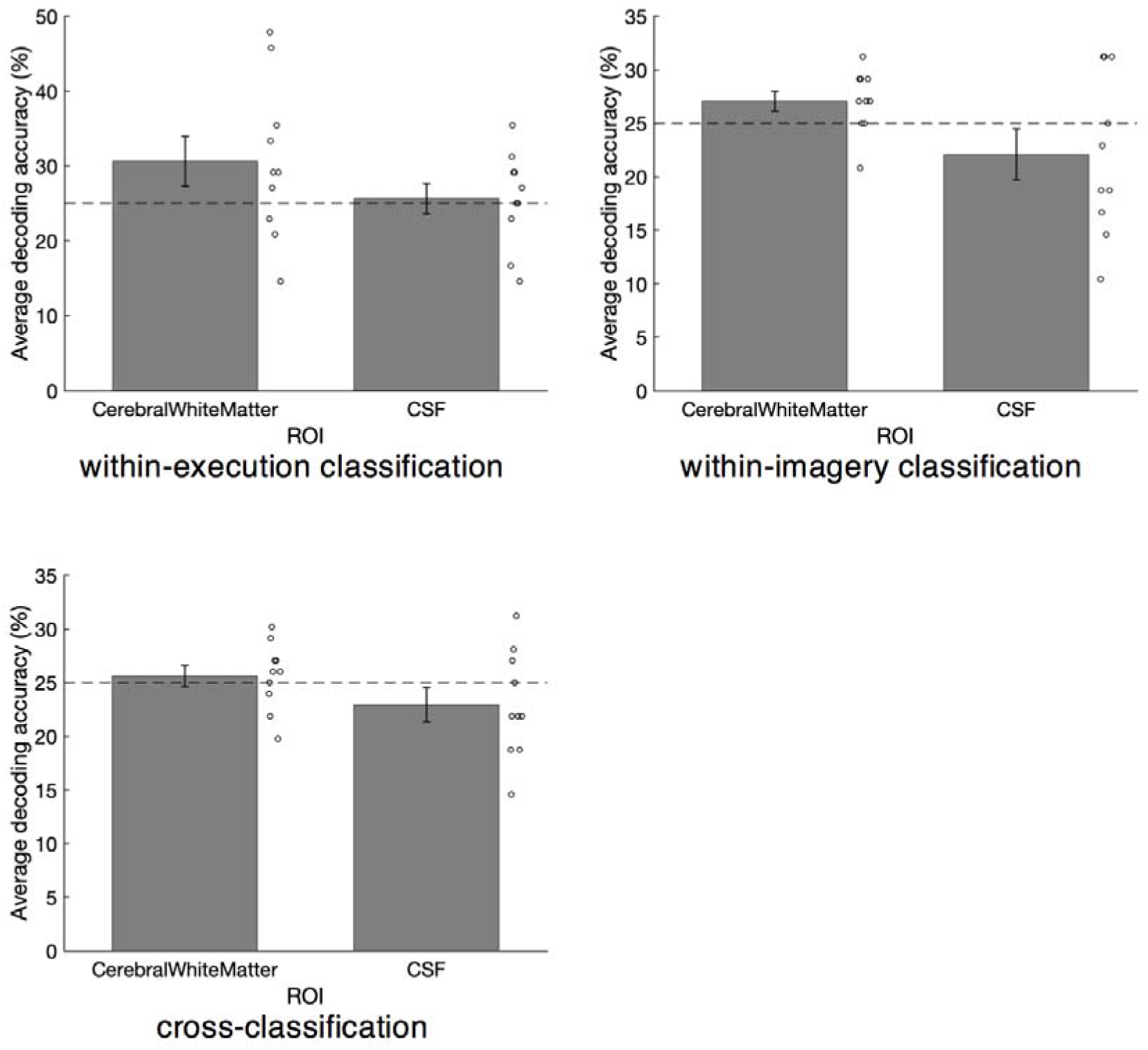
The average decoding accuracy of two control ROIs (cerebral white matter and CSF) in our three main classifications, i.e., within-execution classification (top left), within-imagery classification (top right), and cross-classification (bottom left). Error bars are standard errors. The horizontal dashed line indicates the chance level (25%). Circles indicate single data. In each case, there was no significant decoding accuracy compared to the chance level (|*t*s (9)| < 2.236, *p*s > 0.052 (two-tailed), *d*s < 0.745). Therefore, the significant results of MVPA in the left hand-M1 are unlikely to be by chance.

## References

Amedi, A., Malach, R., Pascual-Leone, A., 2005. Negative BOLD differentiates visual imagery and perception. Neuron 48, 859–872. 10.1016/j.neuron.2005.10.032

Ariani, G., Pruszynski, J.A., Diedrichsen, J., 2022. Motor planning brings human primary somatosensory cortex into action-specific preparatory states. Elife 11, 1–20. 10.7554/eLife.69517

Boccia, M., Sulpizio, V., Bencivenga, F., Guariglia, C., Galati, G., 2021. Neural representations underlying mental imagery as unveiled by representation similarity analysis. Brain Struct. Funct. 226, 1511–1531. 10.1007/s00429-021-02266-z

Decety, J., 1996. The neurophysiological basis of motor imagery. Behav. Brain Res. 10.1016/0166-4328(95)00225-1

Diedrichsen, J., Provost, S., Zareamoghaddam, H., 2016. On the distribution of cross-validated Mahalanobis distances 1–24.

Ejaz, N., Hamada, M., Diedrichsen, J., 2015. Hand use predicts the structure of representations in sensorimotor cortex. Nat. Neurosci. 18, 1034–1040. 10.1038/nn.4038

Fabbri, S., Strnad, L., Caramazza, A., Lingnau, A., 2014. Overlapping representations for grip type and reach direction. Neuroimage 94, 138–146. 10.1016/j.neuroimage.2014.03.017

Faul, F., Erdfelder, E., Buchner, A., Lang, A.G., 2009. Statistical power analyses using G*Power 3.1: Tests for correlation and regression analyses. Behav. Res. Methods 41, 1149–1160. 10.3758/BRM.41.4.1149

Faul, F., Erdfelder, E., Lang, A.G., Buchner, A., 2007. G*Power 3: A flexible statistical power analysis program for the social, behavioral, and biomedical sciences. Behav. Res. Methods 39, 175–191. 10.3758/BF03193146

Grush, R., 2004. The emulation theory of representation: Motor control, imagery, and perception. Behav. Brain Sci. 27, 377–396. 10.1017/S0140525X04000093

Guillot, A., Collet, C., Nguyen, V.A., Malouin, F., Richards, C., Doyon, J., 2009. Brain activity during visual versus kinesthetic imagery: An fMRI study. Hum. Brain Mapp. 30, 2157–2172. 10.1002/hbm.20658

Hall, C.R., Martin, K.A., 1997. Measuring movement imagery abilities: a revision of the movement imagery questionnaire. J. Ment. Imag. 21, 143–154.

Hanakawa, T., 2016. Organizing motor imageries. Neurosci. Res. 104, 56–63. 10.1016/j.neures.2015.11.003

Hanakawa, T., Immisch, I., Toma, K., Dimyan, M.A., Van Gelderen, P., Hallett, M., 2003. Functional properties of brain areas associated with motor execution and imagery. J. Neurophysiol. 89, 989–1002.

Hardwick, R.M., Caspers, S., Eickho, S.B., Swinnen, S.P., 2018. Neuroscience and Biobehavioral Reviews Neural correlates of action : Comparing meta-analyses of imagery, observation, and execution. Neurosci. Biobehav. Rev. 94, 31–44.

Hasegawa, N., 2004. The development of a Japanese version of the revised movement imagery questionnaire. Japanese J. Ment. Imag. 2, 25–34.

Hatta, T., Nakatsuka, Z., 1975. Handedness inventory, Papers on Celebrating 63rd Birthday of Prof. Ohnishi. Osaka City University.

Heming, E.A., Cross, K.P., Takei, T., Cook, D.J., Scott, S.H., 2019. Independent representations of ipsilateral and contralateral limbs in primary motor cortex. Elife 8, 1–26. 10.7554/eLife.48190

Hétu, S., Grégoire, M., Saimpont, A., Coll, M.P., Eugène, F., Michon, P.E., Jackson, P.L., 2013. The neural network of motor imagery: An ALE meta-analysis. Neurosci. Biobehav. Rev. 37, 930–949. 10.1016/j.neubiorev.2013.03.017

Holm, S., 1979. A Simple Sequentially Rejective Multiple Test Procedure. Scand. J. Stat. 6, 65–70.

Jeannerod, M., 1994. The representing brain: Neural correlates of motor intention and imagery. Behav. Brain Sci. 17, 187–202.

Kamitani, Y., Sawahata, Y., 2010. Spatial smoothing hurts localization but not information: Pitfalls for brain mappers. Neuroimage 49, 1949–1952. 10.1016/j.neuroimage.2009.06.040

Kaplan, J.T., Man, K., Greening, S.G., 2015. Multivariate cross-classification: Applying machine learning techniques to characterize abstraction in neural representations. Front. Hum. Neurosci. 9, 1–12. 10.3389/fnhum.2015.00151

Kasess, C.H., Windischberger, C., Cunnington, R., Lanzenberger, R., Pezawas, L., Moser, E., 2008. The suppressive influence of SMA on M1 in motor imagery revealed by fMRI and dynamic causal modeling. Neuroimage 40, 828–837. 10.1016/j.neuroimage.2007.11.040

Kriegeskorte, N., Goebel, R., Bandettini, P., 2006. Information-based functional brain mapping. Proc. Natl. Acad. Sci. 103, 3863–3868. 10.1073/pnas.0600244103

Kriegeskorte, N., Mur, M., Bandettini, P., 2008. Representational similarity analysis-connecting the branches of systems neuroscience. Front. Syst. Neurosci. 2, 4. 10.3389/neuro.06.004.2008

Lebon, F., Horn, U., Domin, M., Lotze, M., 2018. Motor imagery training: Kinesthetic imagery strategy and inferior parietal fMRI activation. Hum. Brain Mapp. 39, 1805–1813. 10.1002/hbm.23956

Ledoit, O., Wolf, M., 2003. Improved estimation of the covariance matrix of stock returns with an application to portfolio selection. J. Empir. Financ. 10, 603–621. 10.1016/S0927-5398(03)00007-0

Lotze, M., Zentgraf, K., 2010. Contribution of the Primary Motor Cortex to Motor Imagery, in: Guillot, A., Collet, C. (Eds.), The Neurophysiological Foundations of Mental and Motor Imagery. Oxford University Press, Oxford, pp. 31–46.

Maldjian, J.A., Laurienti, P.J., Kraft, R.A., Burdette, J.H., 2003. An automated method for neuroanatomic and cytoarchitectonic atlas-based interrogation of fMRI data sets. Neuroimage 19, 1233–1239. 10.1016/S1053-8119(03)00169-1

Monaco, S., Malfatti, G., Culham, J.C., Cattaneo, L., Turella, L., 2020. Decoding motor imagery and action planning in the early visual cortex: Overlapping but distinct neural mechanisms. Neuroimage 218. 10.1016/j.neuroimage.2020.116981

Mur, M., Bandettini, P.A., Kriegeskorte, N., 2009. Revealing representational content with pattern-information fMRI - An introductory guide. Soc. Cogn. Affect. Neurosci. 4, 101–109. 10.1093/scan/nsn044

Naito, E., Kochiyama, T., Kitada, R., Nakamura, S., Matsumura, M., Yonekura, Y., Sadato, N., 2002. Internally simulated movement sensations during motor imagery activate cortical motor areas and the cerebellum. J. Neurosci. 22, 3683–91. 20026282

Ogawa, K., Mitsui, K., Imai, F., Nishida, S., 2019. Long-term training-dependent representation of individual finger movements in the primary motor cortex. Neuroimage 202. 10.1016/j.neuroimage.2019.116051

Oldfield, R.C., 1971. The assessment and analysis of handedness: The Edinburgh inventory. Neuropsychologia 9, 97–113. 10.1016/0028-3932(71)90067-4

Oswald, M.J., Tantirigama, M.L.S., Sonntag, I., Hughes, S.M., Empson, R.M., 2013. Diversity of layer 5 projection neurons in the mouse motor cortex. Front. Cell. Neurosci. 7, 174. 10.3389/fncel.2013.00174

Park, C.H., Chang, W.H., Lee, M., Kwon, G.H., Kim, L., Kim, S.T., Kim, Y.H., 2015. Which motor cortical region best predicts imagined movement? Neuroimage 113, 101–110. 10.1016/j.neuroimage.2015.03.033

Peirce, J.W., 2009. Generating stimuli for neuroscience using PsychoPy. Front. Neuroinform. 2, 1–8. 10.3389/neuro.11.010.2008

Peirce, J.W., 2007. PsychoPy-Psychophysics software in Python. J. Neurosci. Methods 162, 8–13. 10.1016/j.jneumeth.2006.11.017

Persichetti, A.S., Avery, J.A., Huber, L., Merriam, E.P., Martin, A., 2020. Layer-Specific Contributions to Imagined and Executed Hand Movements in Human Primary Motor Cortex. Curr. Biol. 30, 1721–1725.e3. 10.1016/j.cub.2020.02.046

Pilgramm, S., de Haas, B., Helm, F., Zentgraf, K., Stark, R., Munzert, J., Krüger, B., 2016. Motor imagery of hand actions: Decoding the content of motor imagery from brain activity in frontal and parietal motor areas. Hum. Brain Mapp. 37, 81–93. 10.1002/hbm.23015

Porro, C.A., Francescato, M.P., Cettolo, V., Diamond, M.E., Baraldi, P., Zuiani, C., Bazzocchi, M., Di Prampero, P.E., 1996. Primary motor and sensory cortex activation during motor performance and motor imagery: A functional magnetic resonance imaging study. J. Neurosci. 16, 7688–7698. 10.1523/jneurosci.16-23-07688.1996

Roland, P.E., Larsen, B., Lassen, N.A., Skinhoj, E., 1980. Supplementary motor area and other cortical areas in organization of voluntary movements in man. J. Neurophysiol. 43, 118–136. 10.1152/jn.1980.43.1.118

Schnitzler, A., Salenius, S., Salmelin, R., Jousmäki, V., Hari, R., 1997. Involvement of primary motor cortex in motor imagery: a neuromagnetic study. Neuroimage 6, 201–208. 10.1006/nimg.1997.0286

Schroeder, K.E., Irwin, Z.T., Bullard, A.J., Thompson, D.E., Nicole, J., Stacey, W.C., Patil, P.G., Chestek, C.A., 2018. Robust tactile sensory responses in finger area of primate motor cortex relevant to prosthetic control 14, 1–20. 10.1088/1741-2552/aa7329.Robust

Sharma, N., Baron, J.C., 2014. Effects of healthy ageing on activation pattern within the primary motor cortex during movement and motor imagery: An fMRI study. PLoS One 9, 1–8. 10.1371/journal.pone.0088443

Shibasaki, H., Sadato, N., Lyshkow, H., Yonekura, Y., Honda, M., Nagamine, T., Suwazono, S., Magata, Y., Ikeda, A., Miyazaki, M., Fukuyama, H., Asato, R., Konishi, J., 1993. Both primary motor cortex and supplementary motor area play an important role in complex finger movement. Brain 116, 1387–1398. 10.1093/brain/116.6.1387

Stinear, C.M., Byblow, W.D., Steyvers, M., Levin, O., Swinnen, S.P., 2006. Kinesthetic, but not visual, motor imagery modulates corticomotor excitability. Exp. Brain Res. 168, 157–164. 10.1007/s00221-005-0078-y

Tzourio-Mazoyer, N., Landeau, B., Papathanassiou, D., Crivello, F., Etard, O., Delcroix, N., Mazoyer, B., Joliot, M., 2002. Automated anatomical labeling of activations in SPM using a macroscopic anatomical parcellation of the MNI MRI single-subject brain. Neuroimage 15, 273–289. 10.1006/nimg.2001.0978

Zabicki, A., de Haas, B., Zentgraf, K., Stark, R., Munzert, J., Krüger, B., 2017. Imagined and executed actions in the human motor system: Testing neural similarity between execution and imagery of actions with a multivariate approach. Cereb. Cortex 27, 4523–4536. 10.1093/cercor/bhw257

