## Supplementary materials for "The common neural representation in the primary motor area between motor execution and kinesthetic motor imagery"

**EMG recording and processing**

The electromyography (EMG) was measured by BrainAmp ExG MR (BRAIN PRODUCTS) and recorded with the Recorder (BRAIN PRODUCTS) with a sampling rate of 5000 Hz, low cutoff and DC, high cutoff of 1000 Hz. During EMG recording, the EMG clock was synchronized with the clock of an MRI scanner. Offline processing (e.g., noise filtering) of the EMG data was done using Analyzer2 (BRAIN PRODUCTS) in the following order: 1) MR noises were excluded, 2) EMG data were processed using the IIR (zero phase-shift Butterworth) filter with low cutoff: 5 Hz; high cutoff: 100 Hz; Notch: 50 Hz; Bandreject: 17, 33, 50, 67 Hz; bandwidth: 4 Hz, 3) EMG data were downsampled from 5000 Hz to 500 Hz, and 4) EMG data were rectified.

As written in the EMG analysis of results, the analyzed EMG data were extracted from the task blocks of the execution and imagery sessions, and the corresponding timing of the fixation session as a rest period. Generally, some participants can relax well when task onset comes, but others cannot. Cared about such an individual difference, each EMG data extracted from the target block was standardized by the standard deviation of the EMG values that were obtained from the 300 ms period just before each target block.

**Result of the functional localizer**

**Supplementary Table 1:** Anatomical regions, coordinates of peak voxel, and *t*-values of observed activations in the functional localizer of the first study.

| Anatomic region | voxels | MNI coordinates | | | *t*-value |
| --- | --- | --- | --- | --- | --- |
|  |  | x | y | z |  |
| R BA18 | 539 | 6 | -64 | -16 | 15.49 |
| R BA37 |  | 24 | -49 | -25 | 14.63 |
| R BA37 |  | 21 | -55 | -19 | 12.08 |
| L BA4 | 714 | -39 | -22 | 62 | 15.16 |
| L BA3 |  | -33 | -34 | 62 | 13.91 |
| L BA3 |  | -45 | -31 | 59 | 12.69 |
| L BA48 | 181 | -60 | -16 | 11 | 13.39 |
| L BA41 |  | -48 | -31 | 17 | 11.23 |
| L BA48 |  | -54 | -19 | 20 | 7.17 |
| L BA19 | 140 | -27 | -67 | -22 | 10.08 |
| L BA18 |  | -18 | -85 | -16 | 9.02 |
| L BA18 |  | -15 | -76 | -19 | 5.34 |

MNI, Montreal Neurological Institute; L, left hemisphere; BA, Brodmann Area.

**Results of the mass-univariate group analysis (Figure caption)**

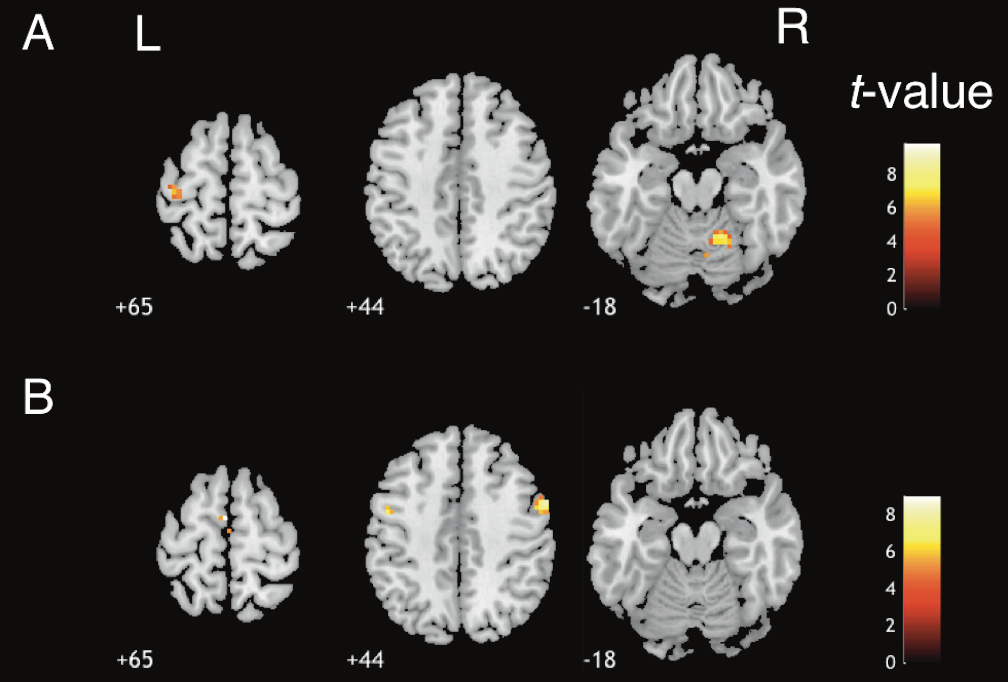

**Supplementary Figure 1:**

(A) The results of mass-univariate group analysis for the execution task session of the first study, with an uncorrected threshold level of *k* ≥ 10 (*p* < 0.0005). The detected left cluster, consisting of BA4 (i.e., X = −39, Y = −19, Z = +65, *t* = 4.862) and BA6, is indicated by the colored area. The detection of BA4 was to be expected, because this region relates to motor control of the contralateral body parts (e.g., Heming et al., 2019). (B) The results of mass-univariate group analysis for the imagery task session in the original study with an uncorrected threshold level of *k* ≥ 10 (*p* < 0.0005). The detected left cluster included BA6 (i.e., X = −3, Y = −4, Z = +65, *t* = 8.981) and is indicated by the colored area, but no voxels from BA4 were detected. SMA is frequently reported as an active region during motor imagery (e.g., Hanakawa, 2016), while M1 activation is rarely reported (Hétu et al., 2013). These results correspond to those in previous studies.

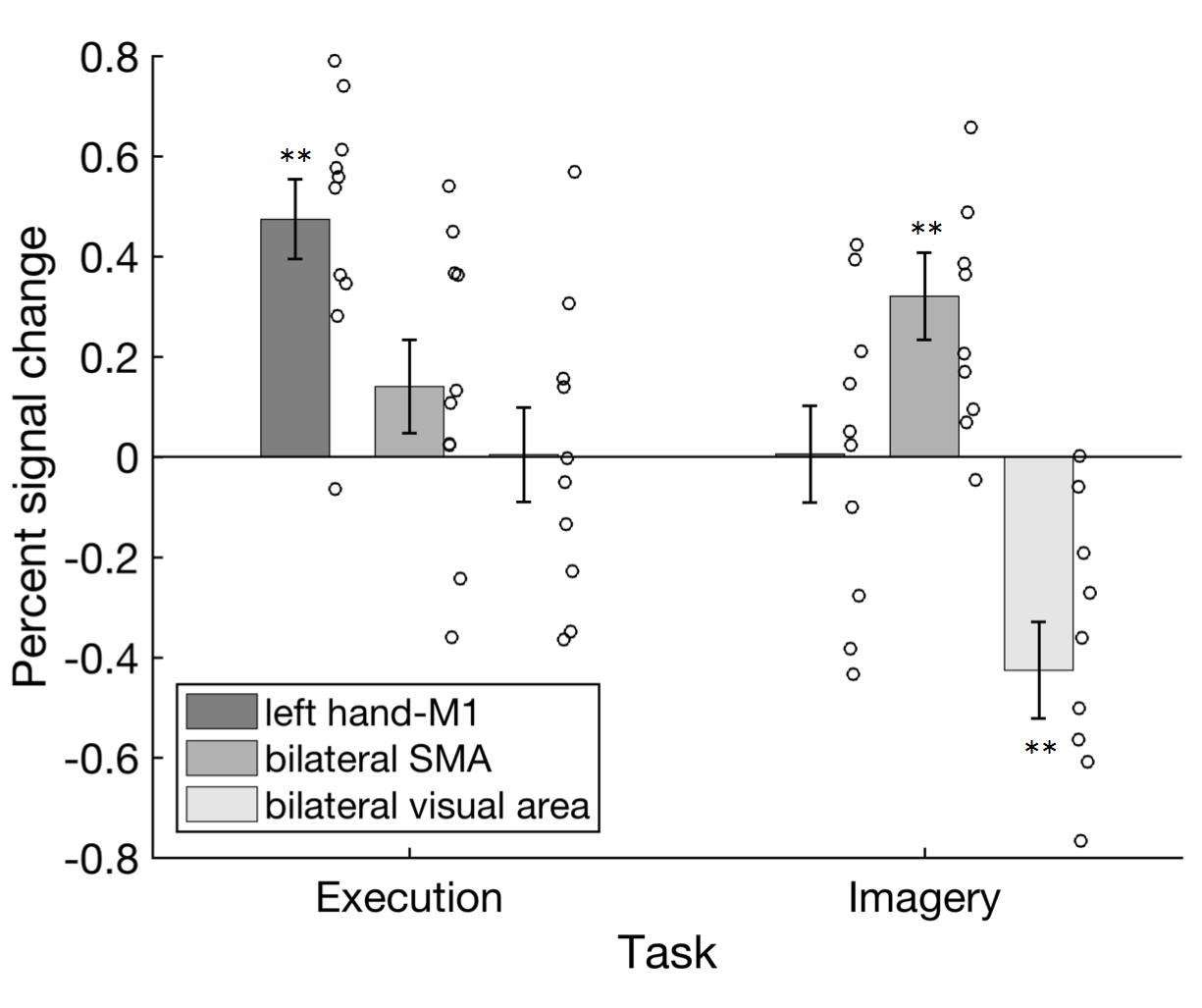

**Supplementary Figure 2:**

The result of the mass-univariate analysis with modeling of rest blocks: The average signal-change rate of each ROI in each task session. Error bars show standard errors. Circles indicate single data. **: *p* < 0.01. Consistent with the result of the mass-univariate analysis without modeling rest, significant differences from rest were found at three locations. Left hand-M1 in motor execution task (*t* (9) = 5.97, *p* < 0.01 (two-tailed), *d* = 1.99), bilateral SMA (*t* (9) = 3.7, *p* < 0.01, *d* = 1.23), and bilateral visual area (*t* (9) = -4.43, *p* < 0.01, *d* = 1.48) in motor imagery task. These results supported our conclusion based on the result of the mass-univariate analysis with modeling rest.

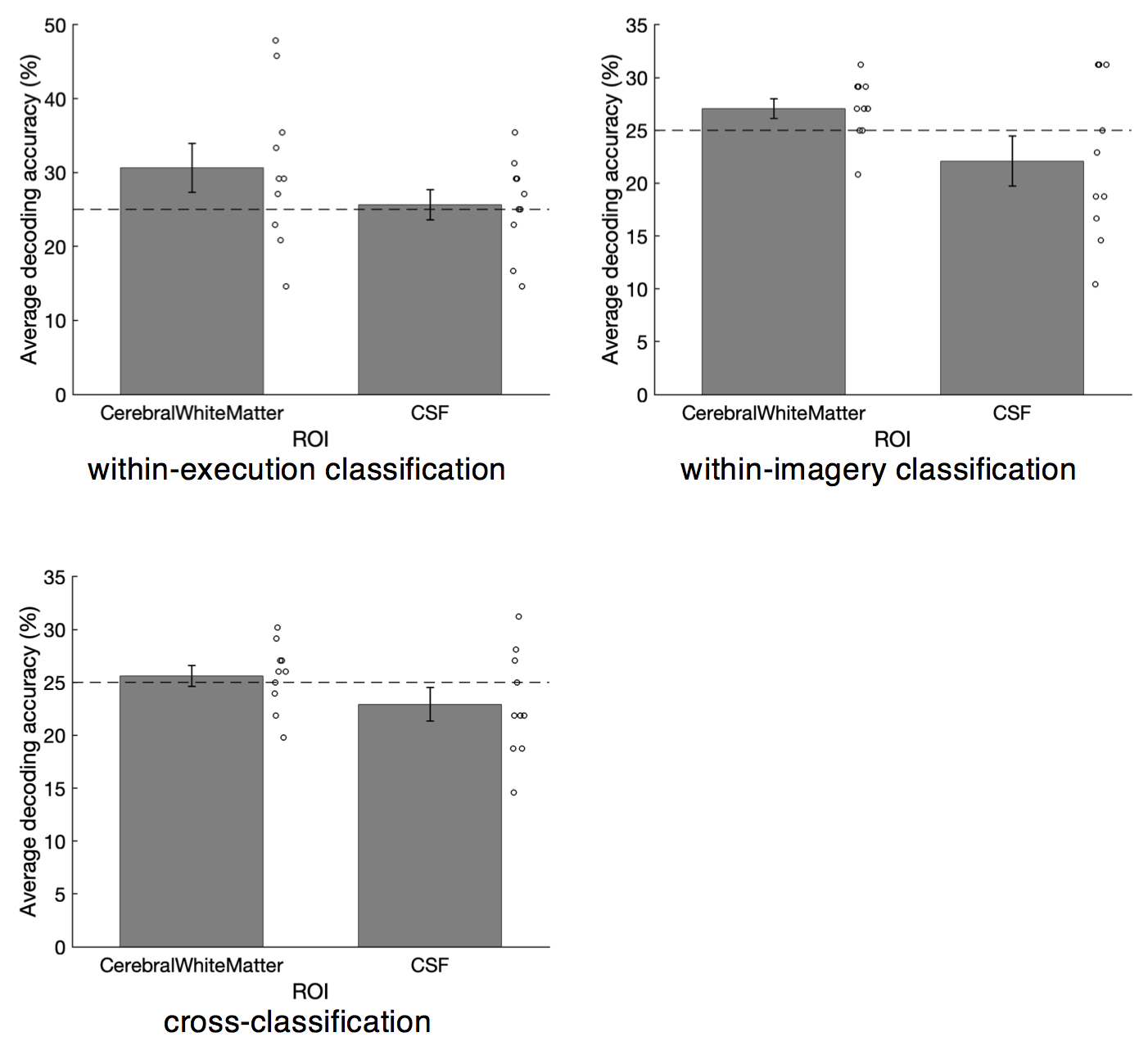

**Supplementary Figure 3:**

The average decoding accuracy of two control ROIs (cerebral white matter and CSF) in our three main classifications, i.e., within-execution classification (top left), within-imagery classification (top right), and cross-classification (bottom left). Error bars are standard errors. The horizontal dashed line indicates the chance level (25%). Circles indicate single data. In each case, there was no significant decoding accuracy compared to the chance level (|*t*s (9)| < 2.236, *p*s > 0.052 (two-tailed), *d*s < 0.745). Therefore, the significant results of MVPA in the left hand-M1 are unlikely to be by chance.
